# Embryonic diversification of adult neural stem cells and ependymal cells

**DOI:** 10.1101/2024.05.12.593751

**Authors:** Shima Yamaguchi, Takaaki Kuniya, Hanae Omiya, Yutaka Suzuki, Masahide Seki, Hideki Ukai, Lingyan Fang, Yujin Harada, Daichi Kawaguchi, Yukiko Gotoh

## Abstract

Both adult neural stem (type B) cells and ependymal (type E) cells in the mouse ventricular-subventricular zone (V-SVZ) are derived from slowly dividing (or quiescent) embryonic neural stem-progenitor cells (NPCs) that are set aside during development. However, it has remained unclear whether fate diversification between adult type B and type E cells actually occurs during embryogenesis. Here we performed single-cell transcriptomic analysis of slowly dividing embryonic NPCs and identified cell subpopulations transcriptionally similar to adult type B or type E cells. The type B-like embryonic cells appeared to emerge before embryonic day (E) 13.5, whereas the type E-like cells became evident between E13.5 and E16.5. Genes differentially expressed in B-like embryonic cells (versus E-like cells) included *Tmem100*, *Cadm2*, and bone morphogenetic protein (BMP)-induced genes. Forced expression of an active form of BMP receptor (ALK2QD), TMEM100, or CADM2 in embryonic NPCs resulted in preferential generation of adult type B cells relative to type E cells in the postnatal brain. Moreover, knockdown of TMEM100 resulted in relative enrichment of type E cells over type B cells. Our results indicate that the embryonic origin of adult type B cells and that of type E cells have already diverged molecularly during embryogenesis, and they have identified key molecular players in this fate bifurcation.

## INTRODUCTION

Embryonic neural stem-progenitor cells (NPCs, also known as radial glial cells) give rise to neurons and glial cells in the central nervous system during development. After completion of brain development and extinction of NPCs in most brain regions, two niches—the ventricular-subventricular zone (V-SVZ) of the lateral ventricle (LV) and the subgranular zone (SGZ) of the hippocampal dentate gyrus—remain to harbor neural stem cells (NSCs, also known as type B cells) that produce neurons and glial cells throughout life in the adult mouse brain ^1–4^. Such newborn neurons are thought to play fundamental roles in circuit plasticity, tissue repair, learning and memory, and innate behaviors, with their dysfunction being implicated in various psychiatric and neurodegenerative diseases ^5–7^. Despite extensive characterization of adult type B cells, however, the ontogeny of these cells—in particular, their relation to embryonic NPCs— has only recently begun to be understood. Whereas adult type B cells in the SGZ are randomly selected from embryonic NPCs in the same region and continue to produce essentially the same cell types (“continuous model”), NPCs that serve as the embryonic origin of adult type B cells in the V-SVZ (also known as preB1 cells) are preserved as a distinct “slowly dividing” population during embryogenesis when other “rapidly dividing” NPCs produce neural cells that contribute to brain development (“set-aside model”) ^3,8–12^. Pulse–labeling with a histone H2B–green fluorescent protein (GFP) fusion protein or with bromodeoxyuridine thus revealed that a subset of embryonic NPCs residing in the ganglionic eminence (GE) experiences a slowing of the cell cycle between embryonic day (E) 13.5 and E15.5 and that a large fraction (>70%) of adult type B cells in the V-SVZ of the LV wall is derived from this slowly dividing (or quiescent) embryonic subpopulation of NPCs ^9,10^.

A key question remains, however, regarding the ontogenic relation between adult type B cells and ependymal cells (hereafter, type E cells). Type E cells line the ventricular wall and play essential roles both in propulsion of cerebrospinal fluid by coordinated beating of their cilia as well as in the support and regulation of adult type B cells ^13^. Adult type B and type E cells are organized into a pinwheel-like structure at the ventricular wall, with the apical endfeet of type B cells being surrounded by a rosette of type E cells ^14^. Of note, precursors of type E cells also manifest a slowing of the cell cycle between E14 and E16 ^15^, and recent studies have uncovered a common precursor of adult type B and type E cells ^16,17^. Pairwise cell fate analyses showed that type E-E and type B-E modes of division were much more frequent than the type B-B mode when the parental cells were labeled at E11.5 to E17.5 and cell fate was determined at postnatal day (P) 10 to P21 ^16,17^. However, given that NPCs that produce only adult type B cells may have already ceased dividing by E13.5 to E17.5, these analyses do not rule out the possibility that embryonic NPCs destined to produce only adult type B cells are also abundant. Furthermore, it has remained unclear whether the differential cell fate apparent after division (that is, E-E versus E-B) has already been determined at the time of division or is decided stochastically (or environmentally) after division during the long maturation process of the cells during postnatal stages ^18,19^. The pursuit of these questions will require determination of whether slowly dividing embryonic NPCs are homogeneous or are actually heterogeneous within the same brain region (that is, irrespective of regional heterogeneity). Furthermore, if these cells are heterogeneous, identification of the molecular basis of this heterogeneity would be expected to provide insight into the mechanisms underlying the choice between adult type B and type E cell fate.

We have now tackled these questions by first performing a single-cell transcriptomic analysis of slowly dividing NPCs in the lateral GE (LGE) of mouse embryos at E16.5. We identified subpopulations of these cells including clusters that show transcriptomic features similar to those of adult type B or type E cells. We further characterized the molecular properties of these adult type B–like and type E–like embryonic NPC clusters and identified their signature genes, which are suggestive of differential cellular status. Finally, manipulation of these signature genes was found to bias NPC fate between type B and type E cells at postnatal stages. Our results suggest that the adult type B and type E lineages are already diverged, at least in part, at embryonic stages.

## RESULTS

### Slowly dividing embryonic NPCs include subpopulations that share transcriptional profiles with adult neural stem (type B) cells or ependymal (type E) cells

To gain insight into the embryonic ontogeny of adult type B and type E cells in the V-SVZ, we performed single-cell (sc) RNA–sequencing (seq) analysis of slowly dividing NPCs—which include the origin of adult type B and type E cells ^9,10,15^—isolated from the dissected LGE and its neighboring neocortical region at E16.5. To facilitate isolation of the slowly dividing cells, we induced transient expression of H2B-GFP in *Rosa-rtTA;TRE-mCMV-H2B-GFP* transgenic mice by injecting a mother with 9-*tert*- butyldoxycycline (9TB-Dox) at E9.5, and we defined label-retaining (slowly dividing) cells (LRCs) as those in the top 10% (H2B-GFP^high^) of GFP fluorescence intensity among CD133^+^CD24^−^ cells prepared from the LGE and its neighboring tissue at E16.5. The scRNA-seq analysis was performed for the H2B-GFP^high^ fraction with the 10x Chromium technology and v2 chemistry platform (Figures 1A and S1A–S1E). We obtained 21 cell clusters that could largely be divided into cells derived from the neocortex (NCX) or the LGE as well as into NPCs (defined by *Hes5* and *Nes* expression) or transit-amplifying cells (TAPs, defined by *Dlx1* or *Dlx2* expression for the LGE and by *Eomes* expression for the NCX) (Figures 1B–1D).

**Figure 1.**
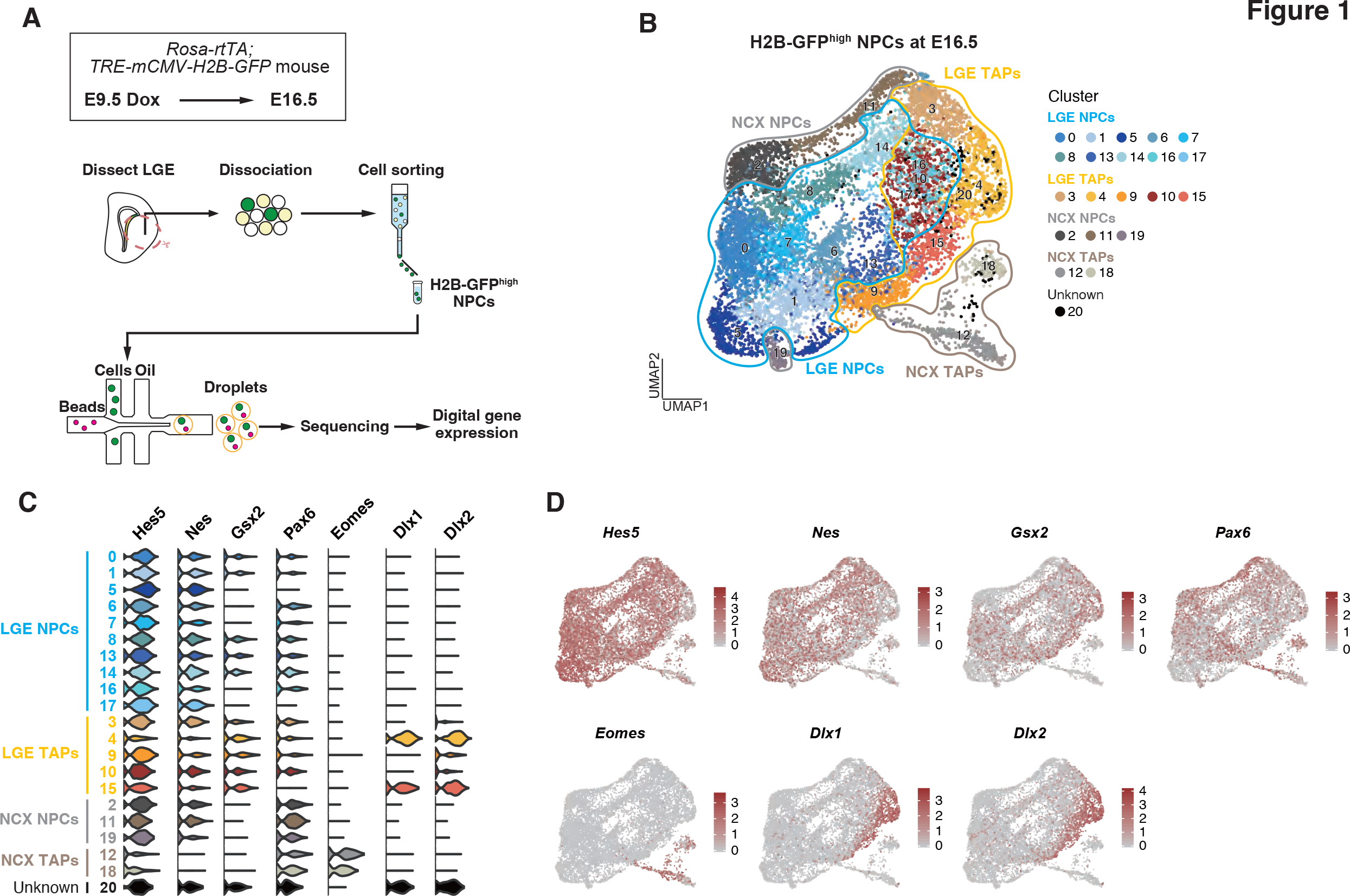
Slowly dividing NPCs have diverse transcriptomic characteristics related to differentiation and brain region (A) Workflow for labeling and single-cell transcriptomic profiling of slowly dividing NPCs. 9TB-Dox (0.25 mg) was injected intraperitoneally into a pregnant *Rosa- rtTA;TRE-mCMV-H2B-GFP* mouse at E9.5, and the LGE and neighboring neocortex (NCX) were dissected from the corresponding embryos at E16.5. NPCs highly retaining the H2B-GFP label were then isolated by fluorescence-activated cell sorting (FACS) and subjected to droplet-based scRNA-seq analysis. (B) Visualization of identified cell clusters by uniform manifold approximation and projection (UMAP). A total of 11,193 cells collected from H2B-GFP^high^ NPCs were classified into 21 clusters. (C) Violin plots for normalized expression of known regional and cell type marker genes. Ventricular zone subregions are defined as follows: NCX as *Pax6*^+^*Gsx2*^−^, and LGE as *Gsx2*^+^. Cell types along the differentiation axis are defined as follows: NPCs as *Hes5*^+^*Nes*^+^, NCX TAPs as *Eomes*^+^, and LGE TAPs as *Dlx1*^+^ or *Dlx2*^+^. (D) Feature plots showing normalized expression of regional and cell type marker genes on the UMAP space. See also Figure S1.

We then investigated the heterogeneity of slowly dividing NPCs in the LGE by performing subcluster analysis for the LGE NPC clusters identified in Figure 1B (Figure 2A). Although these cells were characterized as LRCs, only a subset of the identified clusters was actually in G1 phase of the cell cycle, indicative of a slow cell cycle rather than complete quiescence at this stage (Figure 2B). We then asked whether any of these clusters might be transcriptionally similar to adult type B or type E cells by calculating the sum of the z-scores for the expression of adult type B or type E cell signature genes (designated as the adult type B or type E cell score, respectively) obtained from a reported data set ^20^ (Figure 2D). We found that the adult type B cell score was relatively high in clusters 25, 26, 27, 28, 30, 37, and 40, whereas the adult type E cell score was high in clusters 33 and 22 (Figure 2D). We therefore named the former clusters as “adult type B–like embryonic NPC clusters” (B-like clusters, for short) and the latter clusters as “adult type E–like embryonic NPC clusters I and II” (E-like clusters I and II, for short), respectively. Of note, in addition to adult NSC signature genes such as *Aldoc*, *Cd9*, *Aldh1l1*, and *Slc1a3* ^21^, both *Hey1*, which is required for the long-term maintenance of slowly dividing embryonic NPCs necessary for the generation of adult type B cells ^22^, and *Vcam1*, which is also required for the establishment of adult type B cells during development ^23^, were highly expressed in the B-like clusters (Figures 2C and S2A). Moreover, *Foxj1*, a master regulator of type E cells ^24^, was highly expressed in the E-like clusters (Figure 2C). *Tmem107*, which plays a key role in cilium formation ^25^, was enriched in E-like cluster II, but genes important for type E cell maturation such as *Acta2*, *Ank3*, and *Six3* ^20,26,27^ were not (Figure S2B). These results thus indicated that slowly dividing NPCs in the embryonic LGE include subpopulations that are transcriptionally similar to adult type B or type E cells. E-like cluster I showed a relatively high adult type B cell score, suggesting that this cluster has a transcriptional profile intermediate between those of adult type B and type E cells.

**Figure 2.**
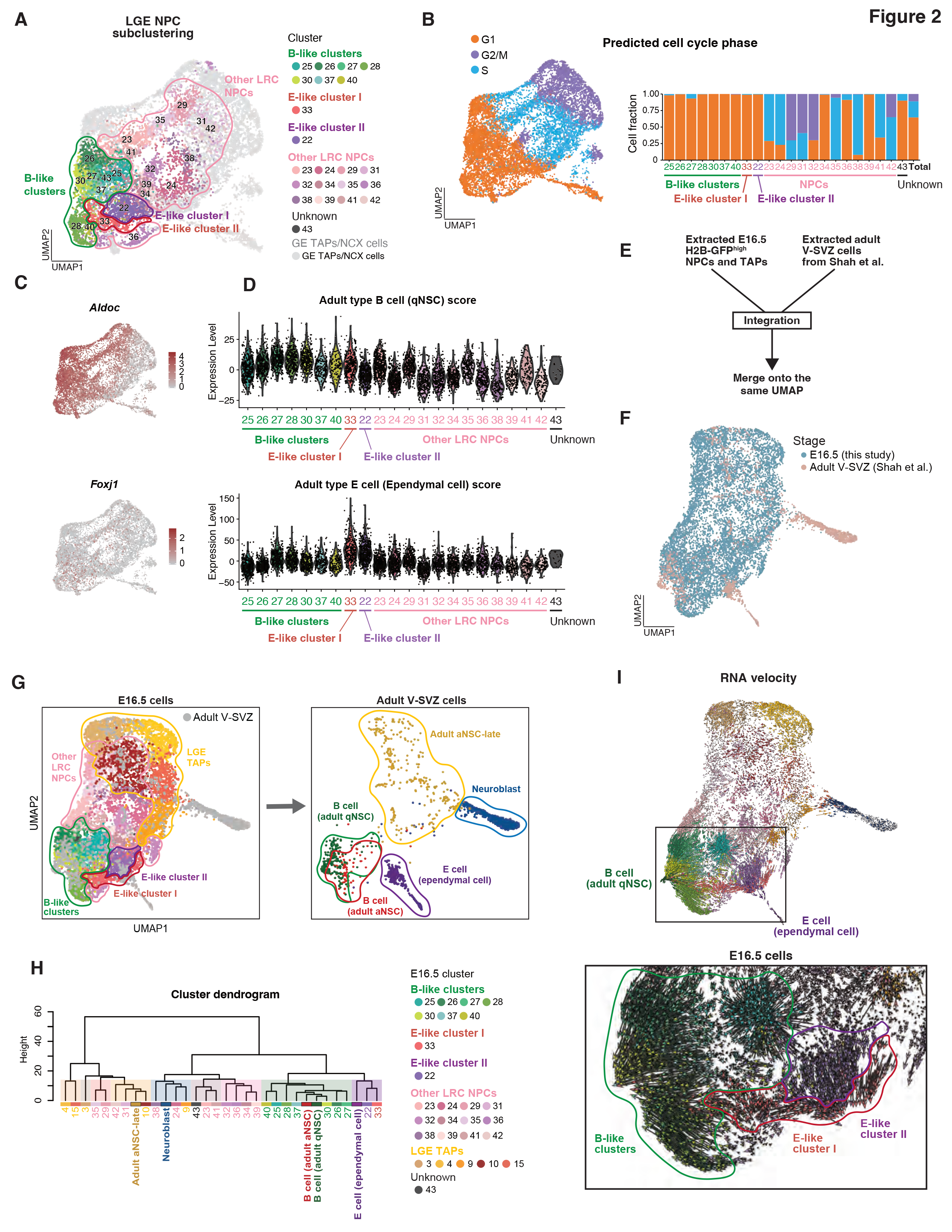
Embryonic B-like cells and E-like cells share transcriptional features with adult type B cells and type E cells (A) Unsupervised graph-based clustering of H2B-GFP^high^ LGE NPCs at E16.5 on the UMAP space. A total of 6093 LGE NPCs was classified into 22 clusters by the subclustering analysis. (B) Feature plot showing cell cycle phase predicted by the *CellCycleScoring* function of Seurat (left), and the proportion of cells in the predicted cell cycle phases for each cluster (right). (C) Normalized expression of *Aldoc* as a quiescent NSC (qNSC) signature gene (top) as well as of *Foxj1* as an ependymal cell signature gene (bottom). (D) Violin plots of the adult type B cell (qNSC) score calculated by summation of the scaled expression of a qNSC signature gene set for each cell (top), as well as of the adult type E (ependymal) cell score calculated by summation of the scaled expression of an ependymal signature gene set for each cell (bottom). (E) Workflow showing integration of E16.5 (this study) and adult V-SVZ (Shah et al., 2018) single-cell data sets. H2B-GFP^high^ NPCs and TAPs of the LGE at E16.5 were integrated with adult V-SVZ neurogenic cells and ependymal cells with the use of Seurat according to a previously described methodology (Butler et al. Nat. Biotechnol., 2018). (F) UMAP plot showing the distribution of E16.5 cells and adult V-SVZ cells. (G) UMAP plots showing cell types among E16.5 cells (left) or adult V-SVZ cells (right). Note that E16.5 B-like clusters map close to adult V-SVZ type B cells, and that E16.5 E-like clusters I and II map close to adult V-SVZ type E cells. (H) The “average PC score” was calculated from the average of principal component (PC) scores of each cell in each cluster for PCs used for UMAP analysis. The cluster dendrogram shows hierarchical clustering of the “average PC score” of each cluster for E16.5 cells and adult V-SVZ cells. E16.5 B-like clusters and E-like clusters I and II are divided into distinct groups and belong to the same groups as adult type B cells and adult type E cells, respectively. (I) UMAP plot for RNA velocity analysis performed for the integrated data as previously described (La Manno et al., Nature, 2018). The starting point of each arrow indicates an individual cell, with the arrow indicating the projected position of the future state based on extrapolated velocity estimates. The boxed region of the upper panel containing E16.5 B-like, E-like cluster I, and E-like cluster II cells is shown at higher magnification in the lower panel. The arrows for E16.5 B-like cells extend toward adult type B cells, those for E16.5 E-like cluster I cells extend toward both adult type B and type E cells, and those for E16.5 E-like cluster II cells extend toward adult type E cells. See also Figure S2.

To examine further the transcriptional similarity between the embryonic B-like and E-like clusters versus adult type B and E cells, we integrated our scRNA-seq data for E16.5 with reported scRNA-seq data for adult V-SVZ cells ^20^ (Figures S2C and S2D) with the use of Seurat v.3.0.1 integration technology that is able to identify pairs of cells in similar biological states. Only LGE NPCs and TAPs from the embryonic data set and quiescent NSCs (qNSCs), activated NSCs (aNSCs), aNSC-late cells (a more mature form of aNSCs), neuroblasts, and type E cells among adult V-SVZ cells were included (Figures 2E and 2F). This integration showed that embryonic B-like clusters mapped to a uniform manifold approximation and projection (UMAP) space overlapping with that occupied by adult qNSCs and aNSCs, that embryonic E-like cluster II mapped to a space close to that occupied by adult type E cells, and that embryonic E-like cluster I mapped to a space located between the positions of adult type B and type E cells (Figure 2G). We then asked whether embryonic B-like and E-like clusters were distinct by performing hierarchical clustering based on the average scores of the principal components used for the UMAP space. This analysis suggested that embryonic B-like and E-like clusters were indeed molecularly segregated (distinguishable), belonging to the same groups as adult qNSCs/aNSCs and E cells, respectively (Figure 2H). In addition, embryonic E-like cluster I was classified into the same group as E-like cluster II and adult type E cells, indicating that the molecular profile of embryonic E-like cluster I was more similar to that of adult type E cells than to that of type B cells. These results together suggested that the diversification between B-like and E-like cells has already occurred, at least to some extent, in mouse embryos.

To investigate whether B-like and E-like embryonic cells can be predicted to become adult type B and type E cells, respectively, we applied RNA velocity analysis ^28^, a computational method for prediction of future transcriptional state, to the multidimensional integrated data set. Embryonic B-like cells were indeed predicted to change their transcriptional state toward that of adult type B cells, whereas embryonic E-like cluster II cells were predicted to change their transcriptional state toward that of adult type E cells (Figure 2I). Embryonic E-like cluster I cells were predicted to change their transcriptional state toward those of both adult type B and type E cells, suggesting that they may have the potential to generate both of these adult cell types. These findings further supported the notion that diversification between adult type B and type E cells actually occurs during embryogenesis.

### Embryonic B-like and E-like NPCs show distinct molecular landscapes and regional distributions

To gain insight into the biological features of B-like and E-like embryonic clusters, we performed gene ontology (GO) analysis for genes that were significantly enriched (*p* < 0.05, nonparametric Wilcoxon rank sum test) in each cluster relative to the remaining cells among LGE NPCs (Figures 3A and 3B). This analysis revealed enrichment of genes related to lipid metabolic processes in embryonic B-like cells, a feature associated with adult quiescent type B cells relative to activated type B cells ^21,29^. Embryonic E-like cluster II showed enrichment of genes related to cilium organization, an essential feature of type E cells, and to regulation of ion homeostasis, a major function of type E cells ^30^. Embryonic E-like cluster I showed enrichment of proliferation-related genes, suggesting that these cells may be at a transition from the mitotic to the quiescent state. Label-retaining NPCs other than B-like and E-like clusters manifested enrichment for mitosis-related genes, indicating that these cells consist mostly of relatively proliferative cells among LRCs (Figure 3B). Together, this analysis suggested that embryonic B-like and E-like NPC clusters already possess certain functional hallmarks of adult type B and type E cells, respectively.

**Figure 3.**
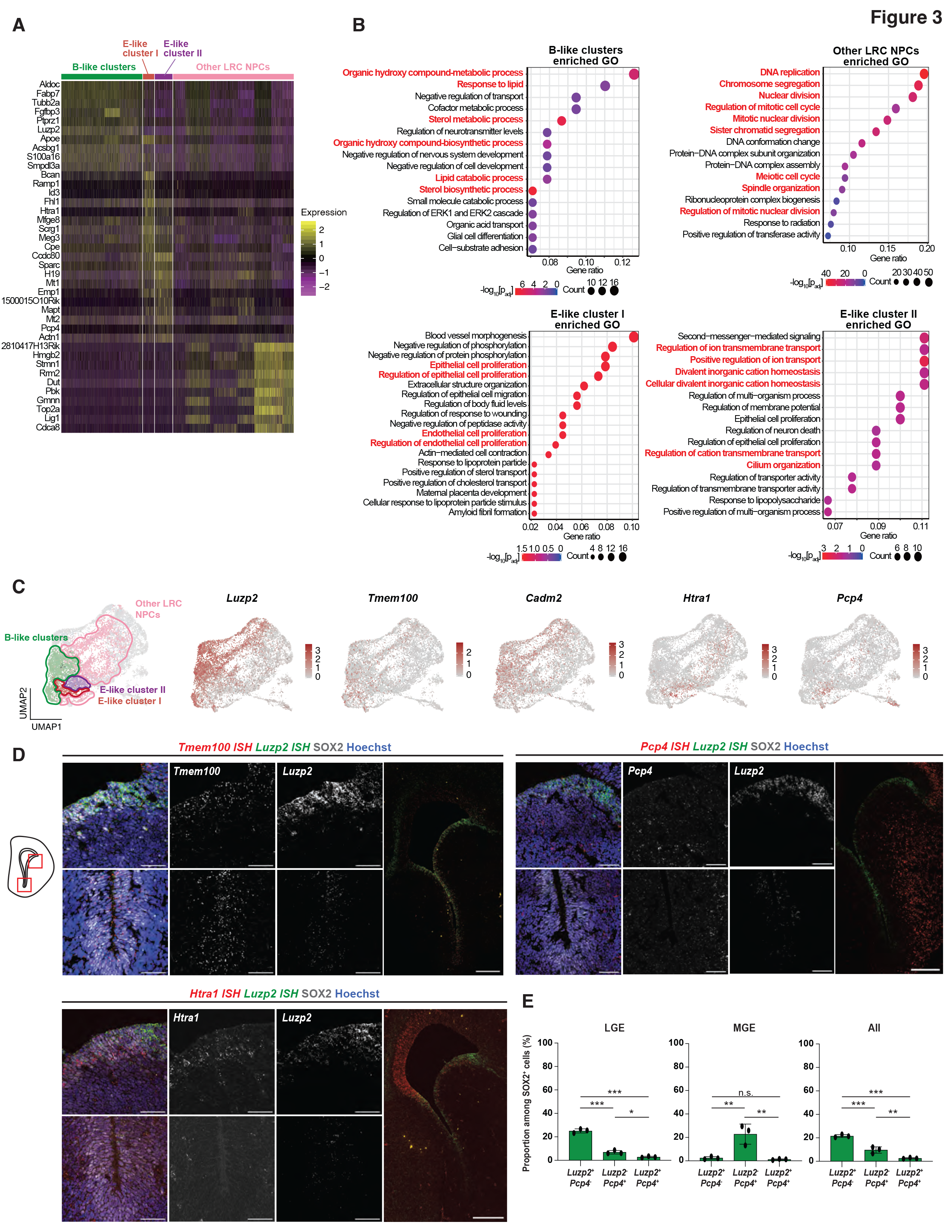
Molecular characterization of B-like and E-like clusters (A) Heat map (scaled by row) showing the expression of signature genes in B-like, E- like I, and E-like II cells as well as in other LRC NPCs at E16.5. Cells are shown in columns and genes in rows. (B) GO term analysis for enriched genes in B-like clusters, E-like cluster I, E-like cluster II, or other LRC NPCs (*p* < 0.05, nonparametric Wilcoxon rank sum test). GO terms related to lipid metabolic processes, cell proliferation or division, ion homeostasis, or cilium organization are highlighted in red. *p*adj, adjusted *p* value. (C) Normalized expression of representative signature genes shown on the UMAP space. *Luzp2*, *Tmem100*, and *Cadm2* are signature genes for B-like cells, whereas *Pcp4* and *Htra1* are signature genes for E-like cells. (D) RNAscope images of *Luzp2* and *Tmem100* (upper left panel), *Luzp2* and *Htra1* (lower left panel), or *Luzp2* and *Pcp4* (upper right panel) in situ hybridization (ISH). Nuclear staining with Hoechst 33342 and immunofluorescence staining of SOX2 are also shown. The rightmost image in each panel shows the entire GE, whereas the smaller upper and lower images are higher magnification views of the dorsal LGE and the MGE, respectively, as indicted in the schematic. Scale bars, 50 µm (higher magnification) or 200 μm (lower magnification). (E) Proportion of *Luzp2*^+^*Pcp4*^−^, *Luzp2*^−^*Pcp4*^+^, or *Luzp2*^+^*Pcp4*^+^ cells among SOX2^+^ cells in the LGE, MGE, or entire GE (all) as determined from images as in (D). Data are means ± SD (*n* = 3 independent experiments). \**p* < 0.05, \*\**p* < 0.01, \*\*\**p* < 0.001; n.s., not significant (two-tailed Student’s *t* test).

Adult type B cells show regional differences in their abundance, molecular features, and progeny cell types within the subdomains of the V-SVZ ^1,2^. Type B cells are enriched in portions of the anterior ventral and posterior dorsal regions of the LV wall, whereas type E cells are enriched in the posterior ventral region ^14,31^. Differences in the expression of type B and type E cell marker proteins (VCAM1 and FOXJ1, respectively) are first apparent after birth in the posterior ventral subdomain of the LV wall ^17^, prompting us to investigate possible regional bias in the abundance of embryonic B-like and E-like cells in the GE. We thus examined the expression patterns of signature genes—*Luzp2*, *Cadm2*, and *Tmem100* for embryonic B-like clusters, and *Htra1* and *Pcp4* for E-like clusters—in the GE at E16.5 by performing RNAscope analysis (Figures 3C and 3D). The expression patterns of *Htra1* and *Luzp2* among SOX2^+^ NPCs in the LGE appeared to be mutually exclusive, with *Htra1^+^* cells and *Luzp2*^+^ cells being located in close proximity (Figure 3D). *Luzp2*^+^ cells and *Tmem100*^+^ cells were distributed throughout the GE, with a marked enrichment in the pallium-subpallium boundary region (Figure 3D). In contrast, *Pcp4*^+^ cells were enriched in the medial GE (MGE). *Luzp2^+^Pcp4^+^*NPCs were significantly less frequent than were *Luzp2^+^Pcp4*^−^ or *Luzp2*^−^*Pcp4^+^* NPCs (Figure 3E), supporting the notion that B-like and E-like embryonic cells are mostly segregated within the GE at E16.5 and that the regional bias of type B and type E cells is already established at this stage.

### Embryonic B-like NPCs become evident by E13.5

We then investigated when B-like and E-like subpopulations emerge in the LGE during embryogenesis by performing scRNA-seq analysis of NPCs and their progeny isolated from the LGE and neighboring neocortical region of wild-type B6J mice at E13.5 (Figures 4A and S1F-S1J). Graph-based clustering of 7039 cells with 2000 highly variable genes and UMAP analysis identified 11 clusters that could be largely divided into NPCs (*Hes5*^+^*Nes*^+^) and TAPs (*Dlx1*^+^ or *Dlx2*^+^ for LGE, *Eomes^+^* for NCX) of the LGE or NCX (Figures 4B and S3A). Subclustering analysis for LGE NPCs from this E13.5 data set identified 10 NPC clusters and 11 committed progenitor clusters based on expression of *Dlx* genes (Figures 4C and S3A). Cluster 27 appeared to represent B-like embryonic NPCs, given its higher adult type B cell score (and high level of expression of the B-like signature genes *Aldoc*, *Luzp2*, and *Cadm2*) relative to other NPC clusters (Figures S3B–S3D). On the other hand, no NPC cluster at E13.5 showed a significantly high adult type E cell score or high expression levels of the E-like signature genes *Foxj1*, *Pcp4*, and *Htra1* (Figures S3B–S3D). Integration of scRNA-seq data sets for NPCs and TAPs of the E13.5 LGE, NPCs and TAPs of the E16.5 LGE, and adult V-SVZ cells revealed that the cells in cluster 27 at E13.5 mapped close to the B-like clusters at E16.5 as well as to adult type B cells, whereas no NPC cluster at E13.5 mapped close to the E-like clusters at E16.5 or adult type E cells (Figures 4D– 4F). RNA velocity analysis of these integrated data indeed showed that the cells in cluster 27 at E13.5 oriented toward adult qNSCs (Figure 4G). Although most LRCs detected at E16.5 are thought to be still proliferating at E13.5 ^9^, about half of the cells in cluster 27 were found to be in G1 phase of the cell cycle (Figure S3E), suggesting that the cell cycle has already slowed in this B-like cluster at E13.5. Consistent with this observation, GO analysis of significantly enriched genes in cluster 27 (B-like cluster) at E13.5 (*p* < 0.05, nonparametric Wilcoxon rank sum test) revealed enrichment of terms related to negative regulation of cell proliferation as well as of terms found to be enriched in B-like clusters at E16.5 including negative regulation of nervous system development, cofactor metabolic process (Figure 4H), cell-substrate adhesion, and sterol biosynthetic process (data not shown), suggesting that establishment of quiescence and B-like characteristics is already under way at E13.5.

**Figure 4.**
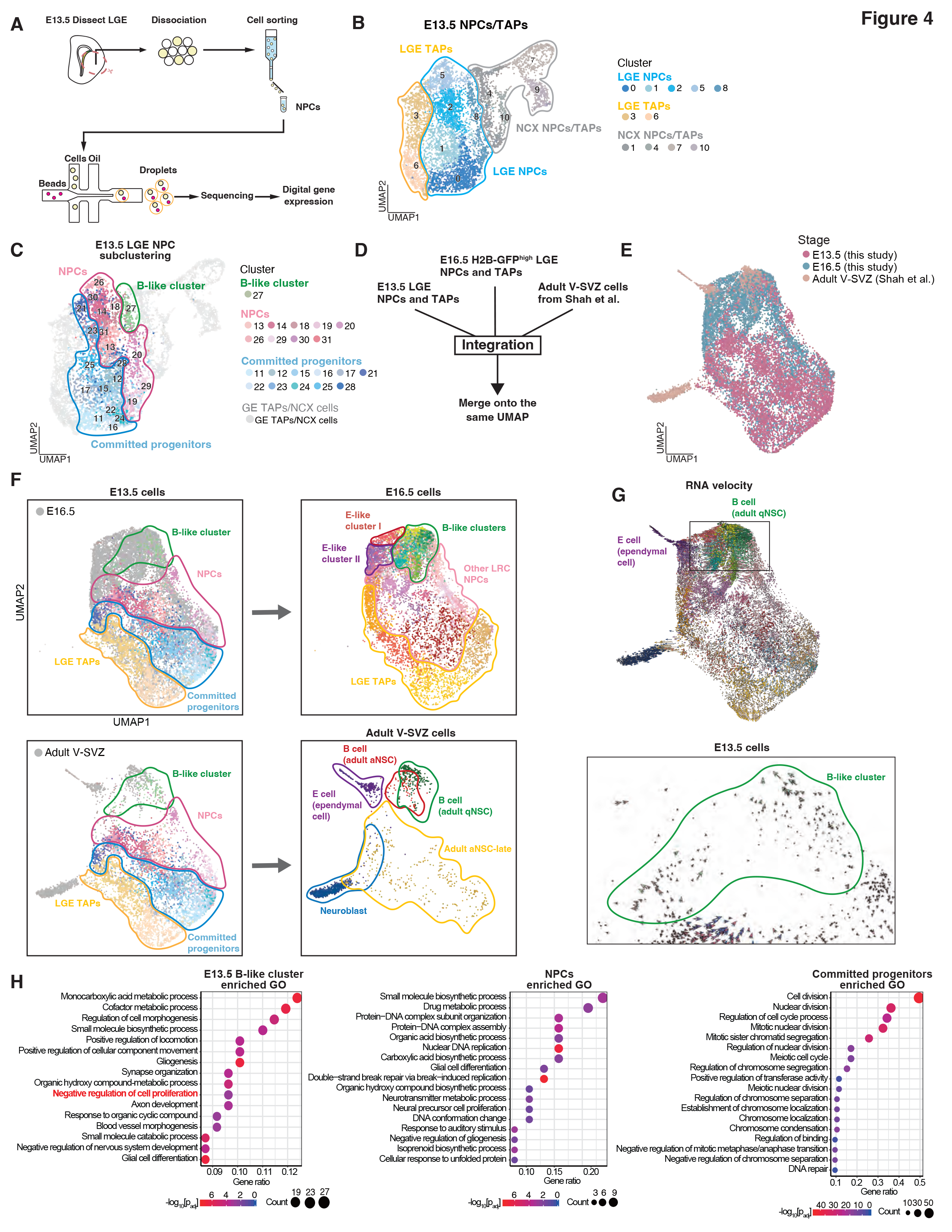
B-like cells are apparent by E13.5, whereas E-like cells are not (A) Workflow for single-cell transcriptomic profiling of E13.5 NPCs. The LGE and neighboring neocortical tissue were dissected from wild-type B6J mouse embryos at E13.5, and NPCs were sorted by FACS and subjected to droplet-based scRNA-seq analysis. (B) Visualization of identified cell clusters by UMAP analysis. A total of 7039 single- cell transcriptomes was classified into 11 clusters. Cell type and brain region for each cluster were defined on the basis of corresponding markers. (C) Unsupervised graph-based subclustering analysis of E13.5 LGE NPCs projected onto the UMAP space. A total of 3903 E13.5 LGE NPCs was classified into 21 clusters. (D) Workflow for integration of E13.5 and E16.5 (this study) as well as adult V-SVZ (Shah et al., 2018) single-cell data sets. E13.5 LGE NPCs and TAPs, E16.5 H2B- GFP^high^ LGE NPCs and TAPs, and adult V-SVZ neurogenic and ependymal cells were integrated with the use of Seurat according to a previously described methodology (Butler et al. Nat. Biotechnol., 2018). (E) UMAP plot showing the distributions of E13.5 cells, E16.5 cells, and adult V-SVZ cells. (F) UMAP plots showing cell types among E13.5 cells, E16.5 cells, and adult V-SVZ cells. E13.5 cell types are plotted in colors with E16.5 cells (upper left) or adult V-SVZ cells (lower left) in gray. E16.5 cell types (upper right) and adult V-SVZ cell types (lower right) are plotted in colors. The E13.5 B-like cluster maps close to both E16.5 B- like clusters and adult V-SVZ type B cells. (G) RNA velocity analysis of the integrated data sets. The boxed region containing E13.5 B-like cells in the upper panel is shown at higher magnification in the lower panel. E13.5 B-like cells extend their arrows toward E16.5 B-like cells and adult V- SVZ type B cells. (H) GO term analysis for enriched genes in the B-like cluster, NPCs, and committed progenitors at E13.5 (*p* < 0.05, nonparametric Wilcoxon rank sum test). The GO term “negative regulation of cell proliferation” is highlighted for the E13.5 B-like cluster. See also Figure S3.

### Activation of BMP signaling in NPCs increases the relative abundance of postnatal type B cells versus type E cells

We then asked whether any of the signature genes or signaling pathways selectively activated in embryonic B-like cells may contribute to NPC fate specification toward adult type B cells (and away from type E cells). We first focused on bone morphogenetic protein (BMP) signaling, given that such signaling promotes adult qNSC maintenance and suppresses type E cell differentiation ^32–36^. Primary NPC cultures (5 days in vitro [DIV]) derived from the V-SVZ at P0 were exposed (or not) to recombinant BMP2 or BMP6 for 1 day and then subjected to RNA-seq analysis (Figures 5A and 5B). Genes upregulated by BMP2 or BMP6 treatment were enriched in E16.5 B-like cells, whereas genes downregulated by BMP2 or BMP6 treatment were enriched in E16.5 E-like cells (Figure 5C). We obtained consistent results for cultures treated with an inhibitor of the BMP type I receptor ALK3 (LDN-193189), given that genes upregulated by LDN-193189 treatment were enriched in E-like cells at E16.5 whereas genes downregulated by LDN-193189 treatment were enriched in E16.5 B-like cells (Figure S4). These results thus showed that genes upregulated by BMP signaling are expressed preferentially in B-like cells compared with E-like cells. We also found that adult type B cell signature genes were enriched in BMP-treated cultures, whereas adult type E cell signature genes were enriched in control cultures (Figure 5D), supporting the notion that activation of BMP signaling skews NPC fate from E-like to B-like, at least in vitro.

**Figure 5.**
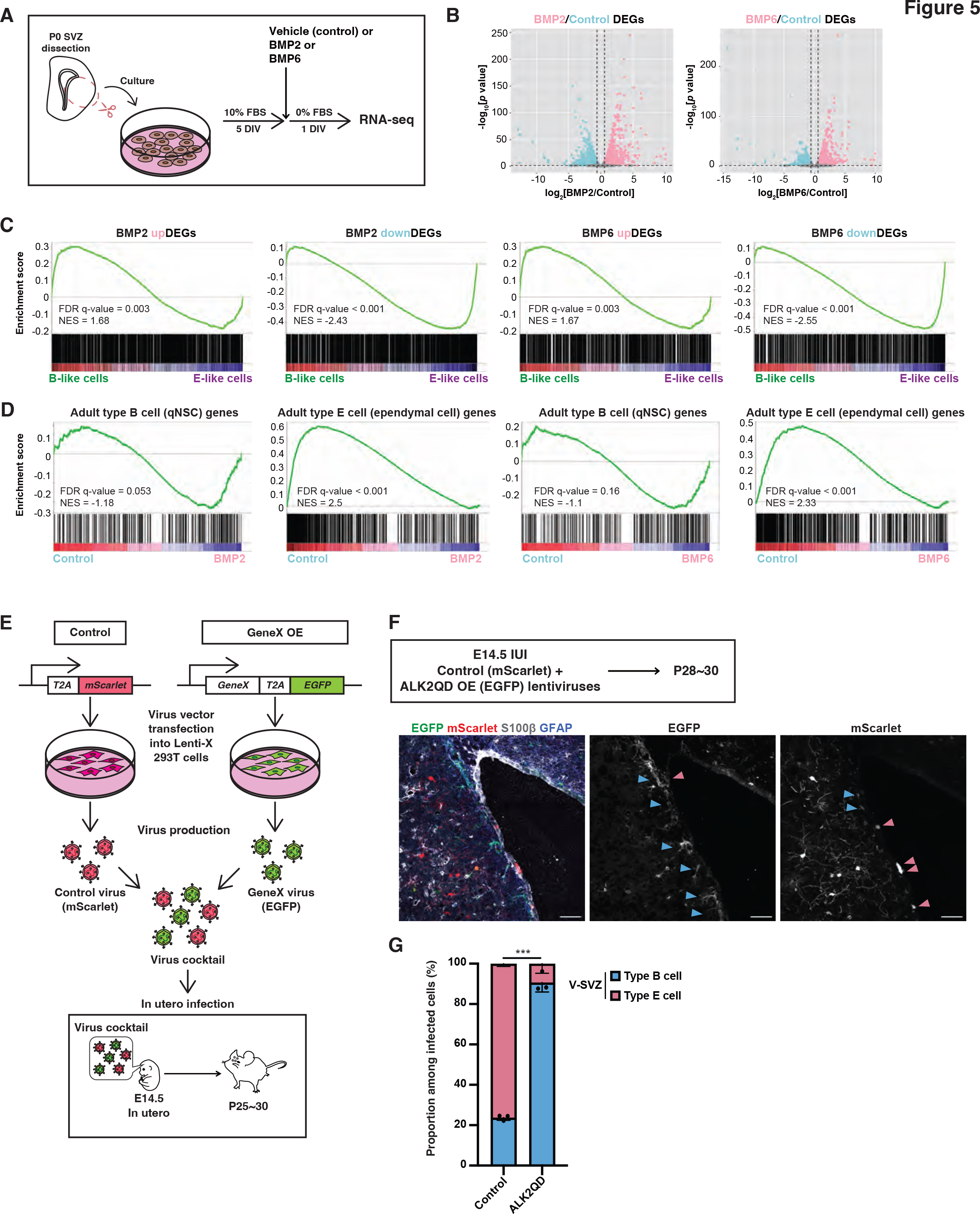
BMP signaling in NPCs increases the ratio of young adult type B cells to type E cells (A) P0 NPC cultures maintained for 5 DIV in the presence of 10% fetal bovine serum (FBS) and then for 1 day without FBS and in the absence or presence of BMP2 (20 ng/ml) or BMP6 (20 ng/ml) were subjected to RNA-seq analysis. (B) Volcano plots showing changes in gene expression induced by BMP2 or BMP6 treatment. Differentially expressed genes (DEGs) that were upregulated by BMP2 or BMP6 treatment (edgeR: *p* < 0.05, log2[fold change] > 0.585) are shown in pink, whereas those downregulated by BMP2 or BMP6 (upregulated in the control) (edgeR: *p* < 0.05, log2[fold change] < –0.585) are shown in light blue. (C) Gene-set enrichment analysis (GSEA) for E16.5 B-like versus E-like cells with DEGs found to be upregulated or downregulated by BMP2 or BMP6 in (B). BMP2- or BMP6-upregulated DEGs were enriched in B-like cells, whereas BMP2- or BMP6- downregulated DEGs were enriched in E-like cells. NES, normalized enrichment score; FDR, false discovery rate. (D) GSEA for BMP2- or BMP6-treated NPC cultures versus control cultures with adult type B cell (qNSC) and adult type E cell gene sets (Shah et al., 2018). Adult type B cell genes were enriched in BMP2- or BMP6-treated cultures compared with control cultures, whereas adult type E cell genes were enriched in control cultures compared with BMP2- or BMP6-treated cultures. (E) Scheme for virus infection experiments. In utero infection (IUI) was performed at E14.5 with the indicated lentivirus cocktail (FU-T2A-mScarlet-W and FU-GeneX-T2A- EGFP-W lentiviruses), and mice were subjected to immunohistofluorescence staining at P25–30. OE, overexpression. (F) The lentiviruses FU-T2A-mScarlet-W and FU-ALK2QD-T2A-EGFP-W were injected into the lateral ventricle of embryos at E14.5, and the resulting mice were subjected to immunofluorescence analysis at P28–30. Brain sections were stained for EGFP, mScarlet, GFAP, and S100β. Light blue and pink arrowheads indicate young adult type B cells and type E cells, respectively. Young adult type B cells were defined as GFAP^+^ cells with apical attachment in the V-SVZ of the lateral wall, and type E cells as S100β^+^ cells. Scale bars, 50 μm. (G) Proportions of young adult type B cells and type E cells among infected cells (control or ALK2QD overexpressing) in the V-SVZ determined from images as in (F). Cells positive for both mScarlet and EGFP were excluded from the analysis. Data are means ± SD (*n* = 3 mice). \*\*\**p* < 0.001, two-tailed Student’s *t* test. See also Figure S4.

To examine whether forced activation of BMP signaling might affect postnatal NPC fate in vivo, we subjected the V-SVZ of wild-type embryos at E14.5 to in utero infection with the lentivirus FU-ALK2QD-T2A-EGFP-W, which confers simultaneous expression of an active mutant (Q207D) of the BMP receptor ALK2 (ALK2QD) and enhanced green fluorescent protein (EGFP), as well as with the control lentivirus FU- T2A-mScarlet-W, which confers expression of monomeric Scarlet (mScarlet) (Figure 5E). We then performed immunohistofluorescence analysis at P28–30 in order to examine the cell types expressing EGFP or mScarlet within the V-SVZ (Figure 5F). We defined type E cells as cells with a high level of S100β expression and a flat shape that aligned with the lateral wall, and type B cells as glial fibrillary acidic protein (GFAP)– expressing cells with apical processes attached to the ventricular surface. Whereas 24 ± 1.3% of control mScarlet^+^ cells became type B cells (and 76 ± 1.3% became type E cells), 91 ± 4.8% of ALK2QD-expressing EGFP^+^ cells became type B cells (and 9 ± 4.8% became type E cells) (Figure 5G). These results indicated that activation of BMP signaling indeed biases the fate of NPCs toward adult type B cells over type E cells.

### Expression of the B-like signature genes *Cadm2* and *Tmem100* increases the relative abundance of postnatal type B cells versus type E cells

We next investigated possible roles of representative B-like signature genes—*Luzp2*, *Cadm2*, and *Tmem100* (Figure 3C)—whose functions in embryonic NPCs and adult NSCs have been unexplored. Permanent labeling (lineage tracing) analysis with lentiviruses (Figure 5E) allowed us to compare the progeny of cells that had been infected with control (FU-T2A-mScarlet-W) or LUZP2-encoding (FU-LUZP2-T2A- EGFP-W) viruses at E14.5 by immunohistofluorescence analysis at P28-30 (Figure S5A). The ratio of type B to E cells within the V-SVZ did not differ significantly between control (mScarlet^+^) and LUZP2-overexpressing (EGFP^+^) cells (Figure S5B), suggesting that the high level of *Luzp2* expression in B-like embryonic clusters does not necessarily affect the fate choice between type B and E cells. On the other hand, comparison of the progeny of cells that had been infected with control (FU-T2A- mScarlet-W) or CADM2-encoding (FU-CADM2-T2A-EGFP-W) viruses at E14.5 revealed that the ratio of type B cells to type E cells within the V-SVZ at P28–30 was greater for the CADM2-overexpressing (EGFP^+^) cells than for control (mScarlet^+^) cells (Figures 6A and 6B). Moreover, the ratio of V-SVZ cells to striatal cells was also greater for CADM2-overexpressing cells than for control cells (Figures 6C and 6D).

**Figure 6.**
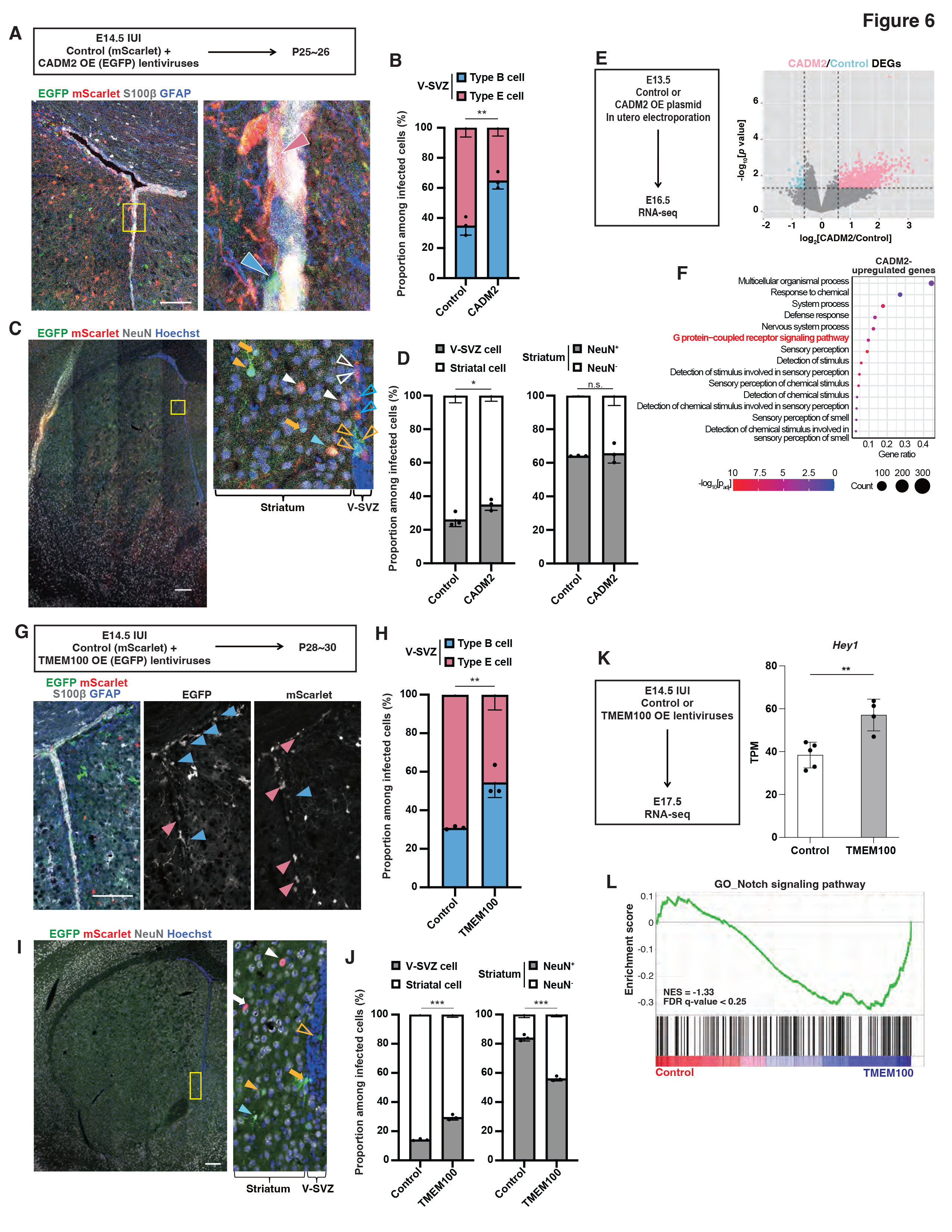
Forced expression of CADM2 or TMEM100 increases the ratio of young adult type B cells to E cells (A) Lentiviruses encoding mScarlet (FU-T2A-mScarlet-W) or both CADM2 and EGFP (FU-CADM2-T2A-EGFP-W) were injected into the embryonic lateral ventricle at E14.5, and brain sections of the resulting mice were subjected to immunohistofluorescence analysis of EGFP, mScarlet, GFAP, and S100β at P25–26. The boxed region in the left panel is shown at higher magnification in the right panel. Light blue and pink arrowheads indicate young adult type B cells (defined as GFAP^+^ cells with apical attachment in the V-SVZ of the lateral wall) and type E cells (defined as S100β^+^ cells), respectively. Scale bar, 100 μm. (B) Proportions of young adult type B cells and type E cells among infected cells (control or CADM2 overexpressing) in the V-SVZ determined from images as in (A). Cells positive for both mScarlet and EGFP were excluded from the analysis. Data are means ± SD (*n* = 3 mice). \*\**p* < 0.01, two-tailed Student’s *t* test. (C) The representative brain section of young adult mouse brains immunostained for EGFP, mScarlet, and NeuN as in (A). Nuclei were stained with Hoechst 33342. The boxed region including the striatum and V-SVZ in the left panel is shown at higher magnification in the right panel. In the striatum, white and yellow closed arrowheads indicate mScarlet^+^NeuN^+^ cells and EGFP^+^NeuN^+^ cells, respectively; yellow closed arrows indicate EGFP ^+^NeuN^−^ cells and the light blue closed arrowhead indicates a cell positive for both mScarlet and EGFP. In the V-SVZ, white and yellow open arrowheads indicate mScarlet^+^ and EGFP^+^ cells, respectively, and blue open arrowheads indicate mScarlet^+^EGFP^+^ cells. Scale bar, 100 μm. (D) Proportions of V-SVZ and striatal cells as well as of NeuN^+^ (neuronal) and NeuN^−^ (nonneuronal) cells in the striatum among corresponding infected cells (control or CADM2 overexpressing) determined from images as in (C). Cells positive for both mScarlet and EGFP were excluded from the analysis. Data are means ± SD (*n* = 3 animals). \**p* < 0.05; n.s., not significant (two-tailed Student’s *t* test). (E) RNA-seq analysis of EGFP^+^CD133^+^CD24^−^ NPCs isolated by FACS from the LGE of embryos at E16.5 that had been subjected to in utero electroporation at E13.5 with a plasmid for EGFP (pCAGEN-EGFP) and either a plasmid for CADM2 (pCAGEN- CADM2) or the corresponding empty plasmid (pCAGEN). The volcano plot shows changes in gene expression induced by CADM2 overexpression. DEGs upregulated by CADM2 (edgeR: *p* < 0.05, log2[fold change] > 0.585) are shown in pink, whereas those downregulated by CADM2 (upregulated in the control) (edgeR: *p* < 0.05, log2[fold change] < –0.585) are shown in light blue. (F) GO enrichment analysis for genes upregulated by CADM2 in (E). (G) Lentiviruses encoding mScarlet (FU-T2A-mScarlet-W) or both TMEM100 and EGFP (FU-TMEM100-T2A-EGFP-W) were injected into the embryonic lateral ventricle at E14.5, and brain sections of the resulting mice were subjected to immunohistofluorescence analysis of EGFP, mScarlet, GFAP, and S100β at P28–30. Light blue and pink arrowheads indicate adult type B cells and type E cells, respectively. Scale bar, 100 μm. (H) Proportions of young adult type B cells and type E cells among infected cells (control or TMEM100 overexpressing) in the V-SVZ determined from images as in (G). Cells positive for both mScarlet and EGFP were excluded from the analysis. Data are means ± SD (*n* = 3 mice). \*\**p* < 0.01, two-tailed Student’s *t* test. (I) The representative brain section of young adult mouse brains immunostained for EGFP, mScarlet, and NeuN as in (G). Nuclei were stained with Hoechst 33342. The boxed region including the striatum and V-SVZ in the left panel is shown at higher magnification in the right panel. In the striatum, white and yellow closed arrowheads indicate mScarlet^+^NeuN^+^ cells and EGFP^+^NeuN^+^ cells, respectively; white and yellow closed arrows indicate mScarlet^+^NeuN^−^ cells and EGFP^+^NeuN^−^ cells, respectively; and the light blue closed arrowhead indicates a cell positive for both mScarlet and EGFP. In the V-SVZ, yellow open arrowheads indicate EGFP^+^ cells. Scale bar, 100 μm. (J) Proportions of V-SVZ and striatal cells as well as of NeuN^+^ (neuronal) and NeuN^−^ (nonneuronal) cells within the striatum among corresponding infected cells (control or TMEM100 overexpressing) determined from images as in (I). Cells positive for both mScarlet and EGFP were excluded from the analysis. Data are means ± SD (*n* = 3 animals). \*\*\**p* < 0.001, two-tailed Student’s *t* test. (K) RNA-seq analysis of EGFP^+^CD133^+^CD24^−^ NPCs isolated by FACS from the LGE of embryos at E17.5 that had been subjected to in utero infection at E14.5 with lentiviruses encoding EGFP (FU-T2A-EGFP-W) or both TMEM100 and EGFP (FU- TMEM100-T2A-EGFP-W). The *Hey1* expression level is shown as transcripts per million (TPM). Data are means ± SD (*n* = 5, 4 animals). \*\**p* < 0.01, two-tailed Student’s *t* test. (L) GSEA of Notch signaling pathway genes for control and TMEM100-overexpressing NPCs analyzed as in (K). Notch signaling pathway genes were enriched in the TMEM100-overexpressing cells. See also Figure S5

These results suggested that CADM2 promotes the maintenance of NPCs and biases the overall fate outcome of NPCs toward adult type B cells rather than type E cells.

CADM2 is an adhesion molecule that suppresses the proliferation of cancer cells ^37^, but its molecular function in NPCs has remained unknown. We therefore performed RNA-seq analysis of EGFP^+^CD133^+^CD24^−^ NPCs isolated at E16.5 after in utero electroporation with an EGFP-encoding vector and with or without a vector for CADM2 at E13.5, and we found that genes upregulated by forced CADM2 expression were enriched for GO terms related to G protein–coupled receptor (GPCR) signaling and stimulus response (Figures 6E and 6F). Given that many GPCR signaling–related genes are also enriched in adult type B cells ^29,38–40^, CADM2 may promote type B cell fate by regulating such signaling pathways.

We then compared the progeny of cells that had been infected with control (FU- T2A-mScarlet-W) or TMEM100-encoding (FU-TMEM100-T2A-EGFP-W) lentiviruses at E14.5 (Figure 6G). The ratio of type B cells to type E cells within the V-SVZ at P28– 30 was much greater for TMEM100-overexpressing (EGFP^+^) cells than for control (mScarlet^+^) cells (Figure 6H). Furthermore, the ratio of V-SVZ cells to striatal cells was also greater for TMEM100-overexpressing cells than for control cells (Figures 6I and 6J). In addition, the ratio of nonneuronal (NeuN^−^) to neuronal (NeuN^+^) cells within the striatum was greater for TMEM100-overexpressing cells than for control cells (Figure 6J). These results indicated that TMEM100 promotes the maintenance of NPCs and biases the overall fate outcome of NPCs toward adult type B cells rather than type E cells, while also biasing the differentiation of striatal cells toward nonneuronal cells (including glial cells) instead of neurons. TMEM100 is a transmembrane protein that is thought to contribute to BMP-mediated endothelial differentiation and other biological processes by modulating various signaling pathways including Notch signaling ^41^. We found that forced activation of BMP signaling by ALK2QD or ALK3QD in embryonic NPCs of the GE increased the abundance of *Tmem100* mRNA (Figures S5C and S5D). In addition, overexpression of TMEM100 in NPCs resulted in enrichment of Notch signaling–related genes (as revealed by GSEA) as well as in increased expression of the Notch target gene *Hey1* (Figures 6K and 6L). Given that *Hey1* plays a key role in the generation of type B cells, TMEM100 may promote this process by regulating the expression of Notch pathway–related genes such as *Hey1* in a manner dependent on BMP signaling.

Finally, we investigated the role of endogenous *Tmem100* in regulation of the generation of postnatal type B and E cells by analyzing the effect of TMEM100 knockdown. We thus performed in utero infection of the V-SVZ at E14.5 with a lentivirus (pLKO-EGFP-*shTmem100*) encoding both a *Tmem100* short hairpin RNA (shRNA) and EGFP as well as with a control lentivirus (pLKO-mScarlet-*shScramble*) encoding both a control (Scramble) shRNA and mScarlet (Figure 7A). The ratio of type B cells to type E cells within the V-SVZ at P28–30 was found to be smaller for TMEM100-depleted (EGFP^+^) cells than for control (mScarlet^+^) cells (Figures 7B and 7C). Neither the ratio of V-SVZ cells to striatal cells nor the ratio of nonneuronal (NeuN^−^) cells to neuronal (NeuN^+^) cells within the striatum differed significantly between TMEM100-depleted cells and control cells (Figures 7D and 7E). These results thus supported the notion that endogenous *Tmem100* biases the overall fate outcome of NPCs toward adult type B cells rather than type E cells.

**Figure 7.**
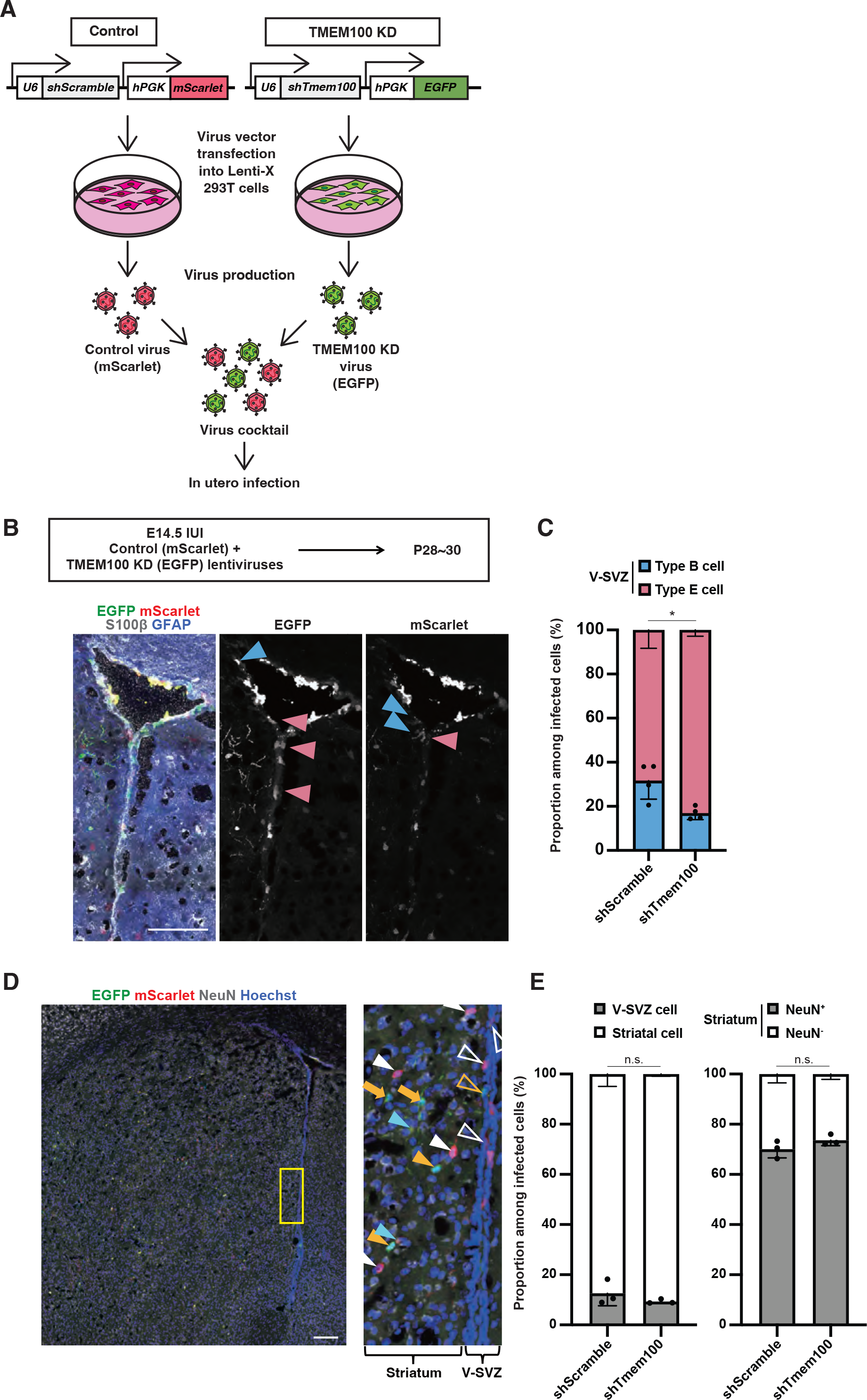
Knockdown of TMEM100 reduces the ratio of young adult type B cells to type E cells (A) Scheme showing the production of lentiviruses for TMEM100 knockdown (KD) experiments. (B) Lentiviruses encoding both mScarlet and a scrambled shRNA (pLKO-mScarlet- *shScramble*) or both EGFP and a TMEM100 shRNA (pLKO-EGFP-*shTmem100*) were injected into the embryonic lateral ventricle at E14.5, and brain sections of the resulting mice were subjected to immunohistofluorescence analysis of EGFP, mScarlet, GFAP, and S100β at P28–30. Light blue and pink arrowheads indicate young adult type B cells (defined as GFAP^+^ cells with apical attachment in the V-SVZ of the lateral wall) and type E cells (defined as S100β^+^ cells), respectively. Scale bar, 100 μm. (C) Proportions of young adult type B cells and type E cells among infected cells (control or TMEM100 depleted) in the V-SVZ determined from images as in (B). Cells positive for both mScarlet and EGFP were excluded from the analysis. Data are means ± SD (*n* = 4 mice). \**p* < 0.05, two-tailed Student’s *t* test. (D) The representative brain section of young adult mouse brains immunostained for EGFP, mScarlet, and NeuN as in (B). Nuclei were stained with Hoechst 33342. The boxed region including the striatum and V-SVZ in the left panel is shown at higher magnification in the right panel. In the striatum, white and yellow closed arrowheads indicate mScarlet^+^NeuN^+^ cells and EGFP^+^NeuN^+^ cells, respectively; yellow closed arrows indicate EGFP ^+^NeuN^−^ cells; light blue closed arrowheads indicate cells positive for both mScarlet and EGFP. In the V-SVZ, white and yellow open arrowheads indicate mScarlet^+^ and EGFP^+^ cells, respectively. Scale bar, 100 μm. Proportions of V-SVZ and striatal cells as well as of NeuN^+^ (neuronal) and NeuN^−^ (nonneuronal) cells within the striatum among corresponding infected cells (control or TMEM100 depleted) determined from images as in (D). Cells positive for both mScarlet and EGFP were excluded from the analysis. Data are means ± SD (*n* = 3 animals). n.s., not significant (two-tailed Student’s *t* test).

## DISCUSSION

Previous studies have shown that common precursors of postnatal type B and type E cells are present in mouse embryos at E11.5–17.5 ^16,17^, but it has remained unclear when and how these precursors become biased toward one fate. We have now identified distinct subpopulations of slowly dividing (label-retaining) NPCs at E16.5 that manifest transcriptional features similar to those of type B or type E cells. We also found that the B-like cells (but not the E-like cells) become distinguishable by E13.5. The maturation process of both type B and type E cells has previously been proposed to occur over a prolonged period and mostly after birth ^19,42^. Our results now indicate that, despite this prolonged maturation process, the fate choice between type B and type E cells may already have taken place around the time of cell cycle slowing (or quiescence onset) at E13.5–E15.5 ^9,10,15^. Our data thus further support the previously proposed notion ^9,10^ that the precursors of adult type B cells in the V-SVZ are “set aside” during embryogenesis as a distinct NPC subpopulation, whereas other rapidly proliferating NPCs contribute to neural development and another quiescent NPC subpopulation contributes to type E cell development. This “set-aside model” of adult type B cell precursors in the V-SVZ contrasts with the “continuous model,” according to which embryonic NPCs (as a whole) gradually become adult type B cells and which has been proposed mainly for adult type B cells that reside in the SGZ of the hippocampal dentate gyrus but also recently for adult type B cells in the V-SVZ ^8,18,19^. It is also of note that interneurons produced by embryonic NPCs in the GE have been shown to establish their transcriptional identity of neuronal subtype after neuronal differentiation rather than before such differentiation (that is, within NPCs) ^43^. Type B cells and type E cells are therefore distinct in that their fate is at least biased and segregated at the NPC stage.

Transcriptional diversification among NPCs during embryogenesis does not necessarily indicate their fate determination. However, if genes or signaling pathways specifically activated in certain NPC clusters promote the establishment of cell fate in a given direction, these NPCs are likely biased toward this fate. Indeed, we found that forced expression of *Tmem100* or *Cadm2* (signature genes highly expressed in the B-like embryonic clusters) in NPCs of the GE from E14.5 increased the ratio of type B cells to type E cells among cells in the V-SVZ at the juvenile postnatal stage (P28–30). We also found that B-like signature genes were enriched together with BMP-induced genes in NPCs, and that forced activation of BMP signaling with the use of an active form of BMP receptor also increased the ratio of type B cells to type E cells among V-SVZ cells. Together, these results suggest that progeny of B-like clusters contribute more to type B cells than to type E cells, although the underlying mechanisms (such as cell proliferation, death, differentiation, and migration) remain to be elucidated.

Moreover, some B-like signature genes identified in this study have been shown to play an essential role in the establishment of postnatal type B cells (such as *Hey1* and *Vcam1*) ^22,23^ as well as in the maintenance or differentiation of type B cells (such as *Ptprz1*, *Apoe*, *Slc1a3*, *Cspg5*, *Cpe*, *Thyh1*, *Id4*, *Slpr1*, *Sepp1*, and *Timp3*) ^34,35,44–54^, further indicative of a preferential contribution of B-like embryonic clusters to type B cells in the adult brain.

We here uncovered previously unappreciated roles of TMEM100 and CADM2 in promoting the generation of postnatal or adult type B cells at the V-SVZ, perhaps at the expense of type E cells. It will be of interest to investigate the functional relation of these proteins to factors that have previously been implicated in the specification of adult type B or type E cells, including DNGR1, VCAM1, Hey1, PRDM16, OLIG2, p53/p21, LAMA5, ANK3, DAG1, GMNN, and GMNC ^16,22,23,26,55–60^. Among these factors, we identified a possible relation between TMEM100 and Notch signaling, which promotes NSC maintenance and suppresses type E cell differentiation. We thus found that overexpression of TMEM100 in NPCs increased the expression of Notch-related genes including that of the Notch target *Hey1*, which plays an essential role in the establishment of postnatal type B cells. Our data also suggest that expression of *Tmem100* promotes the maintenance of embryonic NPCs in the V/SVZ (and limits the number of striatal differentiated cells) while increasing nonneuronal differentiation (and attenuating neuronal differentiation) in the striatum. These effects of TMEM100 are consistent with the reported roles of Notch signaling in promoting NPC maintenance and glial differentiation. It is thus plausible that TMEM100 biases NPC fate toward B-like cells in part through activation of Notch-Hey1 signaling.

BMP signaling may participate in induction of *Tmem100* expression in embryonic B-like clusters, given that BMP signaling has been found to increase such expression in endothelial cells ^41^ and that forced activation of BMP signaling in NPCs in vivo increased *Tmem100* expression in the present study (Figure S5D). BMP signaling has also been implicated in the maintenance of quiescent adult type B cells and in suppression of type E cell differentiation ^32–36^, and we indeed found that forced activation of BMP signaling biased NPC fate toward type B cells rather than type E cells. Of interest, forced expression of active BMP receptors or of TMEM100 in NPCs resulted in marked upregulation of extracellular matrix (ECM)–related genes (Figures S5E–S5G). The possible role of BMP signaling–dependent remodeling of the ECM in establishment of the pinwheel structure characteristic of the adult neurogenic niche in the V-SVZ warrants further investigation. Collectively, these results suggest that BMP signaling biases NPC fate toward type B cells in part through activation of *Tmem100* expression, Notch-Hey1 signaling, and remodeling of the ECM.

In contrast to TMEM100, genes regulated by CADM2 did not appear to overlap with those regulated by BMP or Notch signaling. Moreover, forced expression of CADM2 did not increase the ratio of nonneuronal cells to neuronal cells in the striatum, suggesting that CADM2 functions independently of or in parallel with BMP- TMEM100-Notch signaling. Given that it increased the expression of GPCR-related and stimulus-induced genes, CADM2 might regulate NPC fate through these pathways.

We found that a B-like embryonic cluster, but not an E-like cluster, became transcriptionally evident by E13.5. It is likely that cell cycle arrest is (causally) related to the emergence of B-like clusters but not to that of E-like clusters, given that forced cell cycle arrest in NPCs was previously shown to induce expression of a signature gene set for adult type B cells but not that of signature genes for type E cells (Harada et al. 2021). We indeed here found that the fraction of cells in G1 phase of the cell cycle was already high for the B-like cluster at E13.5 and that the distribution of B-like clusters was similar to that of H2B-GFP label–retaining cells (enriched in the dorsal LGE and ventral GE) ^9^, again supporting an association between cell cycle arrest and the establishment of B-like cells. Cell cycle arrest appears to occur later for type E cells than for type B cells, given the lower level of H2B-GFP retention apparent for type E cells than for type B cells in the postnatal V-SVZ ^9^, which is consistent with our present finding that E-like clusters become transcriptionally evident later than do B-like clusters. The dearth of B-B pairs (relative to E-E and E-B pairs) apparent in pairwise cell fate analyses ^16,17^ is likely attributable to the earlier timing of cell cycle arrest for B-like clusters compared with E-like clusters.

It remains unclear whether E-like clusters emerge from B-like clusters or whether E-like and B-like clusters emerge independently from rapidly dividing NPCs. In the former case, what mechanisms might underlie segregation between B-like and E-like cell fate? Notch signaling may contribute to this fate segregation via the mechanism known as lateral inhibition. In addition, negative feedback mechanisms of BMP signaling (mediated by secreted factors such as Noggin, LRP2, and Chordin-like proteins, for example) may participate in such fate segregation. It would make sense that the embryonic origins of type B and type E cells communicate to develop the pinwheel architecture and to achieve a certain balance between these two cell types that support each other for a lifetime.

## ACKNOWLEDGMENTS

We thank Dr. K Yamamoto (The University of Tokyo) for dieesction faculty; Dr.G Ishii (The University of Tokyo) for FACS; S. Yamaoka (Tokyo Medical and Dental University) for providing us pH2Rwtax lentiviral plasmid; K. Miyazono for providing us BMP receptor constitutively active mutant constructs; the One-Stop Sharing Facility Center for Future Drug Discoveries (The University of Tokyo) for sharing instruments; M. Okajima, R. Yonamine and Y. Kuroda for technical assistance; and members of the Gotoh laboratory for discussion. This research was supported by AMED-CREST and AMED-PRIME of the Japan Agency for Medical Research and Development (JP22gm1310004, JP22gm6110021, JP24gm1310004 for Y.G.), the International Research Center for Neurointelligence (WPI-IRCN) at The University of Tokyo Institutes for Advanced Study (UTIAS), a grant from the Japan Foundation for Applied Enzymology (for T.K.), KAKENHI grants from the Ministry of Education, Culture, Sports, Science, and Technology of Japan and the Japan Society for the Promotion of Science (16J03852, 24K18214 for T.K.; JP16H06279, 22H00431 and 24H02322 for Y.G.).

## AUTHOR CONTRIBUTIONS

S.Y., T.K., and H.O.: conception and design, collection and assembly of data, data analysis and interpretation, and manuscript writing. Y.S. and M.S.: performance of scRNA sequencing experiments. H.U.: technical support Y.H.: data interpretation, funding acquisition, and study supervision. L.F. study supervision:. D.K.: data interpretation, funding acquisition, and study supervision. Y.G.: study conception and design, data interpretation, financial and administrative support, supervision, and manuscript writing. All authors: revision and final approval of the manuscript.

## MATERIALS&METHODS

### Ethics statement

All animals were maintained and studied according to protocols approved by the Animal Care and Use Committee of The University of Tokyo (approval numbers:). All procedures were performed in accordance with the University of Tokyo guidelines for the care and use of laboratory animals.

### Animals

Slc:ICR (ICR) and C57BL/6NCrSlc (B6J) mice were obtained from SLC Japan, and *Rosa26-rtTA* and *TRE-mCMV-H2B-GFP* mice were obtained from The Jackson Laboratory. Mice were maintained at a temperature of 20° to 26°C and relative humidity of 35% to 65% and with a normal 12-h-light, 12-h-dark cycle. They were housed two to six per sterile cage (Innocage, Innovive) containing bedding chips (Palsoft, Oriental Yeast) and were provided with irradiated food (CE2, CLEA Japan) and filtered water ad libitum. Mouse embryos were isolated at various stages, with E0.5 being considered the time of vaginal plug appearance.

### H2B-GFP retention analysis

Expression of the H2B-GFP fusion protein was transiently induced at E9.5 by intraperitoneal injection of 9TB-Dox (0.25 mg) into pregnant *Rosa-rtTA;TRE-mCMV- H2B-GFP* mice, and slowly dividing NPCs were identified as H2B-GFP–retaining NPCs as previously described (Furutachi et al., 2015).

### Tissue preparation for RNA-seq analysis

The LGE and neighboring region of the NCX were rapidly microdissected, and all such tissue from the embryos of each individual pregnant mouse was pooled together. The pooled tissue was dissociated with a papain-based solution (Sumitomo Bakelite or Wako), and the dissociated single cells were stained for 20 min on ice with phycoerythrin- and Cy7-conjugated antibodies to CD133 (1:100 dilution, BioLegend 141210), allophycocyanin-conjugated antibodies to CD24 (1:100, BioLegend 101814) in phosphate-buffered saline (PBS). After that, the cells were stained for 20 min on ice with PE-conjugated streptavidin (1:500 eBioscience 12-4317-87) in PBS. The cells were then immediately subjected to FACS with a FACS Aria instrument (Becton Dickinson). Debris and aggregated cells were removed by gating on the basis of forward and side scatter. NPCs were defined as cells positive for the NPC marker CD133 (also known as prominin) (Weigmann et al., 1997) and negative for the neuronal marker CD24 (Calaora et al., 1996). Gates were set as described previously (Daynac et al., 2013). For isolation of slowly dividing NPCs from *Rosa-rtTA;TRE-mCMV-H2B-GFP* mice treated with 9TB-Dox, we collected NPCs in the top 10% of GFP fluorescence intensity. Data were analyzed with the use of FlowJo software.

### Single-cell library preparation and RNA-seq analysis

Slowly dividing NPCs isolated from *Rosa-rtTA;TRE-mCMV-H2B-GFP* mice at E16.5 and NPCs isolated from wild-type B6J mice at E13.5 were used to prepare single-cell sequencing libraries according to the 10x Genomics Chromium Single Cell 3’ Reagent Guidelines (v2 chemistry). Sequencing was performed with an Illumina Hiseq 2500 instrument (read 1, 26 cycles; index, 8 cycles; read 2, 98 cycles, paired end). Reads were processed with the 10x Genomics Cell Ranger (version 2.0.0) pipeline. FASTQ files generated from Illumina sequencing output were mapped to the reference mouse genome (mm10) with the STAR algorithm. The average read depth was 10,136 and 17,128 reads per cell for the slowly dividing NPC (E16.5) and wild-type NPC (E13.5) libraries, respectively.

### scRNA-seq analysis

#### Quality control, cell clustering, and visualization

The R toolkit Seurat was used for quality control and downstream analysis of scRNA-seq experiments (Macosko et al., 2015). All functions were run with default parameters, unless otherwise specified. Before analysis, we removed predicted blood cells, immune cells, oligodendrocyte progenitor cells (OPCs), and mature neurons as well as MGE cells on the basis of detection of the following genes: *Adm*, *Cldn5*, *Higd1b*, *Igfbp7*, *Mgp*, *Pglyrp1*, *Rgs5*, *Slc38a5*, and *Vtn* for blood cells; *C1qa*, *C1qb*, *C1qc*, *Ccl3*, *Ccl4*, *Ctss*, and *Il1b* for immune cells; *Olig1* for OPCs; *Sst* for mature neurons; and *Nkx2-1* for MGE cells. The raw expression matrix generated by Cell Ranger was imported into Seurat (v.3.0.1) in R (v.3.6.1). The distributions of unique molecular identifier (UMI) counts and genes for each cell in each data set were plotted (Figures S1B and S1G). For generation of Seurat objects, genes expressed in <3 cells and cells with <200 genes were excluded. Low-quality cells were filtered on the basis of the numbers of UMI counts and overall genes as well as the proportion of mitochondrial genes per cell; cells with <200 or >4000 genes and with >5% of genes mapped to the mitochondrial genome were discarded for slowly dividing NPCs at E16.5 (Figure S1C), and those with <200 or >4800 genes and >5% of mitochondrial genes were removed for wild-type NPCs at E13.5 (Figure S1H).

After exclusion of low-quality cells, gene expression was normalized and highly variable genes (top 2000 genes) were identified according to the Seurat tutorial. Scaling of the data was performed with all detected genes. Principal component (PC) analysis was conducted with highly variable genes for identification of PCs appropriate for application to clustering and UMAP visualization by JackStraw analysis (Figures S1D and S1I). PC1 to PC16 and PC1 to PC15 were used for analysis of the E16.5 and E13.5 data sets, respectively. For subclustering analysis, we reanalyzed highly variable genes and performed scaling and JackStraw analysis again, resulting in the use of PC1 to PC12 and PC1 to PC11 for the subclustering of E16.5 and E13.5 data sets, respectively. Graph-based clustering and UMAP visualization were performed with the selected PCs. For UMAP visualization, we selected “umap-learn” as the umap-method and “correlation” as the metric in the *RunUMAP* function. For cell clustering, we chose resolutions of 1.1 for the E16.5 data set and 0.6 for the E13.5 data set; a resolution of 2 was used for the subclustering of both E16.5 and E13.5 data. Each cluster was then annotated on the basis of the expression of selected marker genes. Total gene and UMI counts were plotted for each cell in each subcluster (Figures S1E and S1J). All UMAP plots, gene expression patterns visualized on UMAP plots, violin plots, and heat maps were generated with Seurat.

#### Cell cycle scores

A cell cycle score was calculated with the *CellCycleScoring* function of Seurat (v.3.0.1) run with default parameters.

#### Cell type scores

To explore the transcriptional similarity between clusters identified from our experiments and adult type B or type E cells, we calculated the sum of the z-scores for the expression of adult type B (qNSC) or type E cell signature genes obtained from a previous data set (Shah et al. 2018) (GEO: GSE100320). We designated the resulting scores as adult type B and type E cell scores, respectively.

#### Integration of developmental and adult data sets

To compare transcriptional profiles of embryonic stages obtained from our experiments with that of the adult stage, we used the publicly available data set of Shah et al. (GEO: GSE100320). We first reanalyzed the adult data set with Seurat (v.3.0.1) and identified each cluster mentioned in the previous study (Figures S2C and S2D). We then extracted cells with transcriptional profiles of NPCs and TAPs in the LGE from our embryonic data sets, and type B cells (both qNSCs and aNSCs), type E cells, aNSC-late cells, and neuroblasts from the adult data set for integration analysis. We integrated the data sets with the use of the *FindIntegrationAnchors* and *IntegrateData* functions, and then performed scaling for the integrated data sets. After PC analysis and Jackstraw analysis, we chose PC1 to PC14 and PC16 for the integration of E16.5 and adult data sets, and PC1 to PC19 for that of E13.5, E16.5, and adult data sets. Each cluster defined in each data set was visualized on the UMAP space for the integrated data sets.

In addition, for comparison of the transcriptional profiles of our embryonic clusters with those of the adult clusters, we performed hierarchical clustering by Ward’s method with the average PC scores used in the UMAP space for each cluster. On the basis of these results, we calculated the sum of the squares of the distances between each cluster and the mean for the group to which it belongs, with the whole being divided into *k* groups. We defined *k* as the number of groups for which the sum of the squares was minimized, and colored over the branches of the hierarchical clustering (Figure 2H).

#### Identification of DEGs and GO analysis

To identify DEGs for each cluster, we used the *FindAllMarkers* function of Seurat (v.3.0.1) and defined genes with an adjusted *p* value of <0.05 as signature genes. For analysis of the biological functions of such genes, we performed GO analysis with the *enrichGO* function of R package clusterProfiler (v.3.14.3), with “ont” (ontology) set to “BP” (biological process), “pvalueCutoff” to 0.05, “pAdjustMethod” (method for *p* value adjustment) to “BH” (Benjamini-Hochberg), “universe” (background genes) to all genes detected in the data set, and “qvalueCutoff” to 0.1. The output was processed with the *simplify* function from clusterProfiler and visualized as dot plots.

#### RNA velocity analysis

RNA velocity analysis was performed with velocyto.py (v.0.17.17) in Python3 and velocyto.R (v.0.6), Seurat (v.4.3.0), and SeuratWrappers (v.0.3.1) in R (v.4.1.2), with all functions being run with default parameters, unless otherwise specified. Estimation of unspliced and spliced mRNAs was computed with the following parameters in velocyto.py: *velocyto run10x* SAMPLEFOLDER GTFFILE (“refdata-cellranger- mm10-1.2.0” obtained from 10x Genomics Cell Ranger). The output files were converted to Seurat objects in R and processed by the *SCTransform* function. RNA velocity was calculated with the *RunVelocity* function of SeuratWrappers and with the following parameters: deltaT = 1, kCells = 25, and fit.quantile = 0.02. The calculated data for our embryonic data sets were integrated with those for the adult data set (Shah et al. 2018) with the use of the *FindIntegrationAnchors* and *IntegrateData* functions. Each velocity was visualized on the UMAP space with the use of the *show.velocity.on.embedding.cor* function and with the parameter “neighborhood cell numbers” set to 600. Arrows for each cell were plotted with velocyto.py and the parameter “quiver_scale” set to 50.

### Immunofluorescence analysis

For immunohistofluorescence staining of coronal brain sections, mice were anesthetized by intraperitoneal injection of pentobarbital (Nacalai Tesque) and then transcardially perfused with ice-cold 4% paraformaldehyde (Merck) in PBS. The brain was then removed, exposed to the same fixative overnight at 4°C, equilibrated with 30% (w/v) sucrose in PBS, embedded in OCT compound (Tissue TEK), and frozen. Coronal cryosections (thickness of 12–16 μm) were exposed to Tris-buffered saline containing 0.1% Triton X-100 and 2% donkey serum (blocking buffer) for 2 h at room temperature, incubated first overnight at 4°C with primary antibodies in blocking buffer and then for 2 h at room temperature with Alexa Fluor–conjugated secondary antibodies (Thermo Fisher Scientific) in blocking buffer, and mounted in Mowiol (Calbiochem).

Primary antibodies for immunostaining included chicken anti-GFP (1:1000 dilution, Abcam ab13970), rat anti-GFP (1:1000, Nacalai Tesque GF090R), anti-SOX2 (1:200, Cell Signaling Technology 3728), anti–red fluorescent protein (1:1000, MBL PM005), anti–red fluorescent protein (1:500, Rockland 600-401-379), mouse anti- S100β (1:200, Sigma-Aldrich S2657), rabbit anti-S100β (1:500, Abcam ab52642), anti-GFAP (1:1000, Abcam ab4674), and anti-NeuN (1:500, Biolegend 834501). Nuclei were stained with Hoechst 33342 (1:1000, Molecular Probes).

Fluorescence images were obtained with a laser confocal microscope (Leica TCS-SP5 or Zeiss LSM 880) and were processed with the use of LAS AF (Leica), ZEN (Zeiss), Photoshop CS (Adobe), and Image J (U.S. National Institutes of Health) software.

### RNAscope analysis

RNAscope analysis was performed with the use of an RNAscope 2.5 HD Assay-RED kit (ACD). In brief, OCT compound-embedded sections prepared from fresh tissue were depleted of paraffin, rehydrated, exposed to hydrogen peroxide, and processed for target retrieval and protease treatment. *Luzp2*, *Tmem100*, *Pcp4*, and *Htra1* probes (ACD) were applied for hybridization. After signal amplification, stained slides were imaged with a Zeiss LSM 880 microscope.

### Lentivirus preparation

Concentrated lentiviruses were produced by transfection of Lenti-X 293T cells (6.0 × 10^6^ per 100-mm dish) with lentiviral vectors (pCAG-VSVG, psPAX2, pAdvantage, pH2Rwtax), each forced expression vector or knock down vector with the use of TransIT-293 Transfection Reagent (Mirus). Conditioned medium collected 48 to 72 h after the onset of transfection was cleared of debris by low-speed centrifugation (1000 × *g*, 5 min), passed through a 0.45-μm filter, and centrifuged for 120 min at 20°C first at 50,000 × *g* and then at 26,000 × *g*. The final virus pellet was suspended in PBS.

### In utero infection and electroporation

Introduction of lentivirus or plasmid DNA into NPCs of developing mouse embryos was performed as previously described (Furutachi et al. 2015). Concentrated virus suspension or plasmids (2 μg/μl) were injected into the LV. For electroporation, electrodes were positioned at the flanking ventricular regions of each mouse embryo, and four pulses of 45 V for 50 ms were applied at intervals of 950 ms with the use of an electroporator (CUY21E, Tokiwa Science). The uterine horn was then returned to the abdominal cavity to allow the embryos to continue to develop.

### Plasmid vectors

Complementary DNAs for mouse proteins were used for all forced expression experiments. Vectors used for lentivirus production included pCAG-VSVG, psPAX2, pAdvantage, pH2Rwtax (Yamaoka et al. 1996), FUGW, FU-2A-mScarlet-W, FU- Tmem100-2A-EGFP-W, FU-ALK2QD-2A-EGFP-W, FU-ALK3QD-2A-EGFP-W, FU-LUZP2-2A-EGFP-W, FU-CADM2-2A-EGFP-W, pLKO-mScarlet-*shScramble*, and pLKO-EGFP-*shTmem100*. Plasmid vectors used for electroporation were pCAGEN- EGFP and pCAGEN-CADM2.

### Postnatal NPC culture

The LV wall was dissected from ICR mice at P0, and dissociated cells prepared therefrom were transferred to 24-well plates coated with poly-D-lysine (Sigma-Aldrich) at a density of 1 × 10^6^ cells/ml in a proliferation medium consisting of Dulbecco’s modified Eagle’s medium containing high glucose (DMEM-High Glucose 4.5, Gibco) and supplemented with 10% FBS (Gibco) and 1% penicillin-streptomycin (Invitrogen). After 5 DIV, the medium was switched to a differentiation medium (proliferation medium without FBS).

Recombinant human/mouse/rat BMP2 or mouse BMP6 (R&D Systems) was added to cultures at 50 ng/ml in differentiation medium after incubation of the cells for 5 DIV in proliferation medium. BMP2 and BMP6 were dissolved in a solution containing 4 mM HCl and 0.1% bovine serum albumin, which was also used as a vehicle control. Alternatively, after culture for 5 DIV in proliferation medium, the cells were exposed to LDN-193189 (Sigma-Aldrich) at 0.1 μM in proliferation medium.

LDN-193189 was dissolved in distilled water.

### Bulk RNA-seq analysis

Cells positive for CD133 and EGFP and negative for CD24 were isolated by FACS from the LGE of E17.5 embryos that had been subjected to in utero infection at E14.5 with a lentivirus for EGFP alone (control) or for EGFP together with TMEM100 or ALK2QD, or ALK3QD, or from the LGE of E16.5 embryos that had been subjected to in utero electroporation at E13.5 with a plasmid for EGFP alone (control) or together with a plasmid for CADM2. In addition, postnatal NPCs cultured with BMP2, BMP6, or LDN-193189 were collected 1 day after treatment onset.

Purified RNA was used for library construction for RNA-seq analysis. Template preparation was performed with a SMART-Seq Stranded Kit (Takara Bio), and the constructed templates were subjected to deep sequencing analysis with the HiSeq X Ten platform (Illumina) to obtain 150-bp paired-end reads. About 40 million sequences were obtained for each RNA-seq analysis. Sequence reads were mapped to the reference mouse genome (mm10) with the use of Hisat2. Only uniquely mapped and “deduplicated” reads with no base mismatches were used. Reads were processed by TMM (weighted trimmed mean of M-values) normalization as implemented in the R package “edgeR.” DEG analysis for representative samples was also performed with edgeR. GO enrichment analysis was conducted with gProfiler software ^61^.

### GSEA

Normalized gene expression data for bulk RNA-seq analysis were subjected to GSEA. Enrichment of signature genes was assessed with GSEA software. The gene sets used for GSEA are shown in Supplementary Data Table S1.

### Statistical analysis

Data are presented as means ± SD as indicated, and they were compared with the two-tailed Student’s *t* test. A *p* value of <0.05 was considered statistically significant.

## SUPPLEMENTAL FIGURE LEGENDS

**Figure S1.**
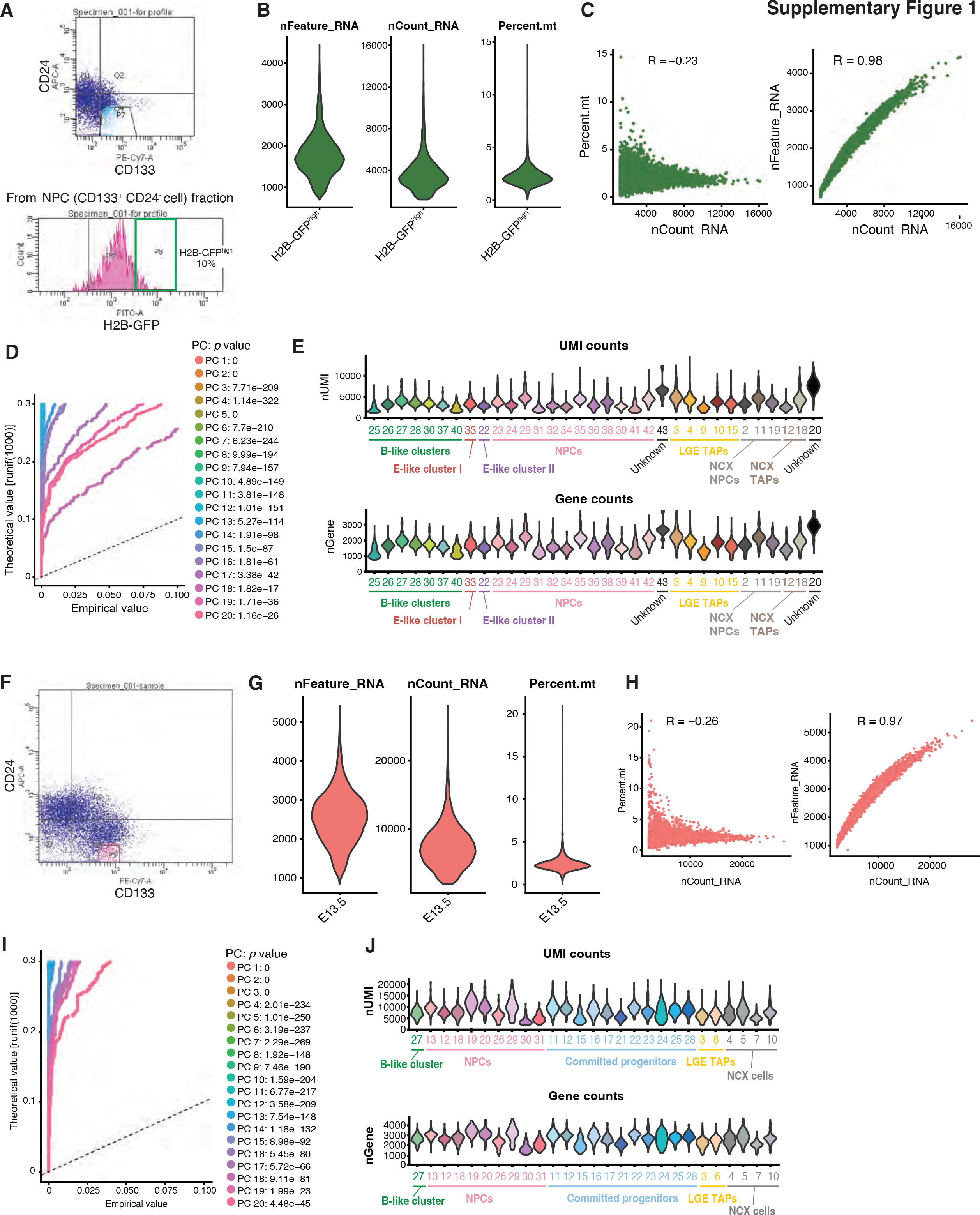
Quality control for scRNA-seq analysis of slowly dividing NPCs at E16.5 and of NPCs at E13.5 (A) Flow cytometry gating scheme for NPCs of the LGE of *Rosa-rtTA;TRE-mCMV- H2B-GFP* mice at E16.5 that highly retain the H2B-GFP label (H2B-GFP^high^ NPCs). Gating for the NPC fraction defined as CD133^+^CD24^−^ cells (top 7–9%) is shown in the upper panel, and that for slowly dividing cells (top 10% of H2B-GFP fluorescence intensity) is shown in the lower panel. (B) Violin plots of UMIs (unique molecular identifiers), gene counts, and the percentage of mitochondrial gene counts for individual cells among slowly dividing cells at E16.5. nFeature_RNA indicates the number of unique genes, nCount_RNA indicates the total number of UMIs, and percent.mt indicates the percentage of reads that map to the mitochondrial genome. (C) Pearson correlation analysis for UMI counts (nCount_RNA) and either the percentage of mitochondrial gene counts (percent.mt) or total gene counts (nFeature_RNA) for individual cells. (D) Jackstraw plot for determination of principal components (PCs) to use for dimension reduction and clustering. PC1 to PC16 were identified as PCs with strong enrichment and a low *p* value. (E) Violin plots of UMI counts (top) and gene counts (bottom) for each E16.5 cluster. (F) Flow cytometry gating scheme for the NPC fraction defined as CD133^+^CD24^−^ cells (top 7–9%) from the LGE of wild-type mice at E13.5. (G) Violin plots of UMIs, gene counts, and the percentage of mitochondrial gene counts for individual NPCs at E13.5. (H) Pearson correlation analysis for UMI counts and either the percentage of mitochondrial gene counts or total gene counts for individual cells. (I) Jackstraw plot used for determination of PCs to use for dimension reduction and clustering. PC1 to PC15 were identified as PCs with strong enrichment and a low *p* value. (J) Violin plots of UMI counts (top) and gene counts (bottom) for each E13.5 cluster.

**Figure S2.**
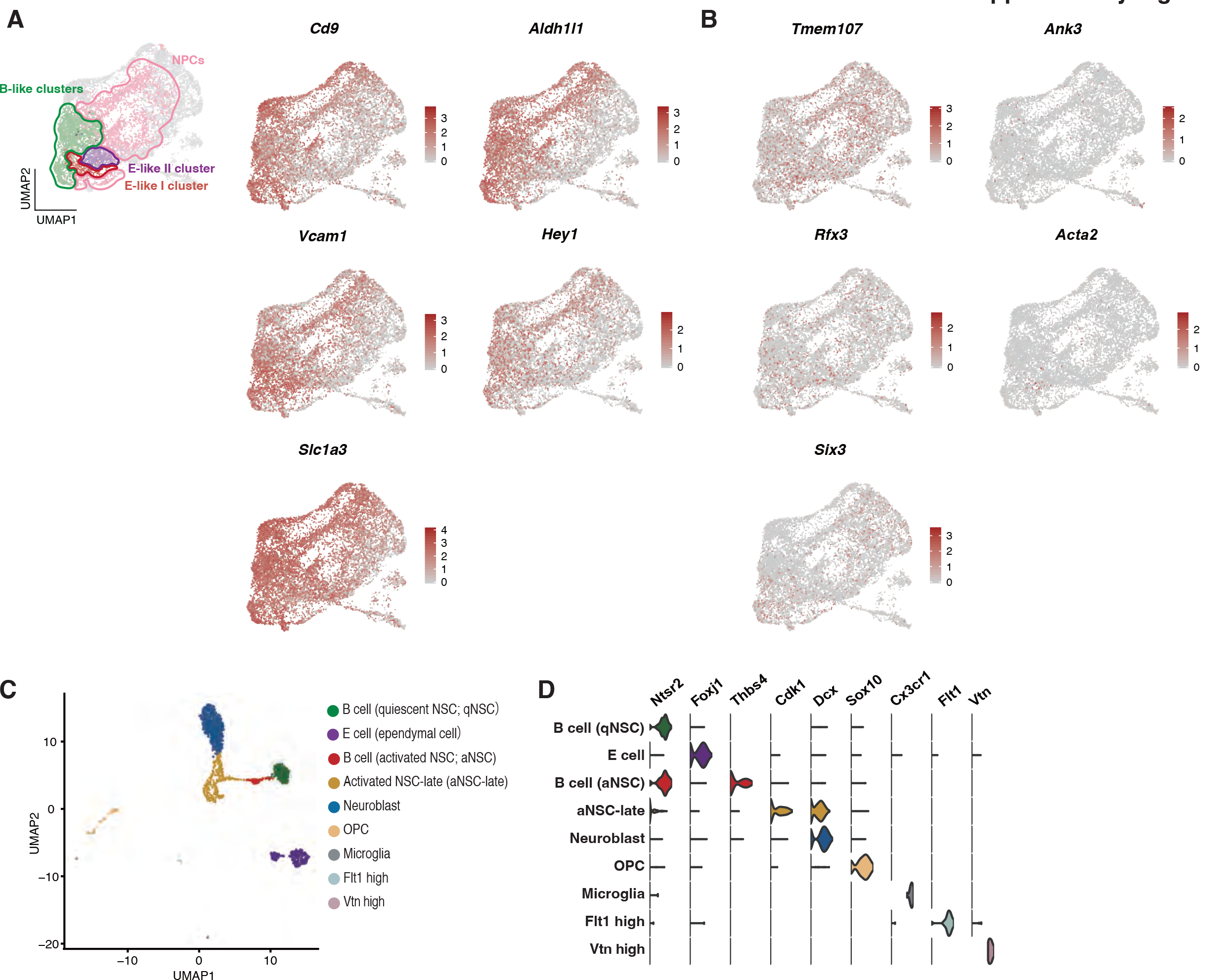
Cluster analysis of slowly dividing E16.5 NPCs and adult V-SVZ cells (A) Normalized expression of adult type B cell signature genes (*Cd9*, *Aldh1l1*, *Slc1a3*, *Hey1*, and *Vcam1*) in slowly dividing E16.5 NPCs shown on the UMAP space. (B) Normalized expression of adult type E (ependymal) cell signature genes (*Tmem107*, *Rfx3*, *Acta2*, *Ank3*, and *Six3*) in slowly dividing E16.5 NPCs shown on the UMAP space. (C) Reanalysis of published scRNA-seq data for adult V-SVZ cells (Shah et al.) with the use of Seurat v.3.0.1. Cluster annotation is as in the previous study. OPC, oligodendrocyte progenitor cell. (D) Violin plots for normalized expression of marker genes for each cluster used in the previous study.

**Figure S3.**
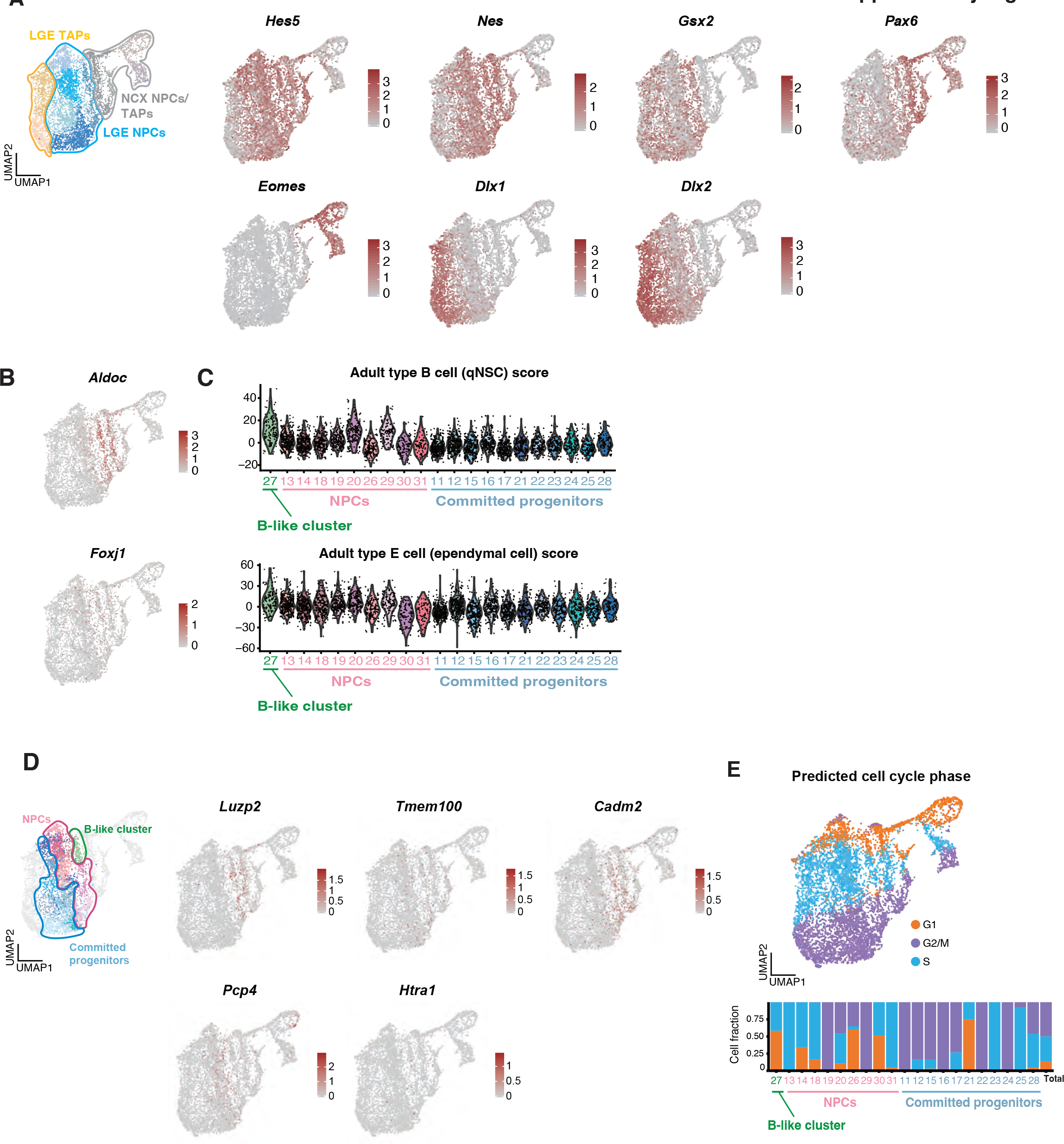
Cluster analysis of E13.5 NPCs (A) Feature plots showing normalized expression of regional and cell type marker genes on the UMAP space for NPCs isolated from the LGE and neighboring NCX of wild-type mice at E13.5. Ventricular zone subregions are defined as follows: NCX as *Pax6^+^Gsx2*^−^ and LGE as *Gsx2*^+^. Cell types along the differentiation axis are defined as follows: NPCs as *Hes5*^+^*Nes*^+^, NCX TAPs as *Eomes*^+^, and LGE TAPs as *Dlx1*^+^ or *Dlx2*^+^. (B) Normalized expression of *Aldoc* as an adult type B cell (qNSC) signature gene (top) and of *Foxj1* as an adult type E cell signature gene (bottom) shown on the UMAP space. (C) Violin plots of the adult type B cell (qNSC) score (top) and the adult type E (ependymal) cell score (bottom) calculated by summation of scaled expression of corresponding signature gene sets (Shah et al). (D) Normalized expression of representative signature genes on the UMAP space. *Luzp2*, *Tmem100*, and *Cadm2* are signature genes for B-like cells, whereas *Pcp4* and *Htra1* are signature genes for E-like cells. (E) Feature plot showing cell cycle phase predicted by the *CellCycleScoring* function in Seurat (top), and the proportion of cells in each predicted cell cycle phase for each cluster of E13.5 LGE NPCs.

**Figure S4.**
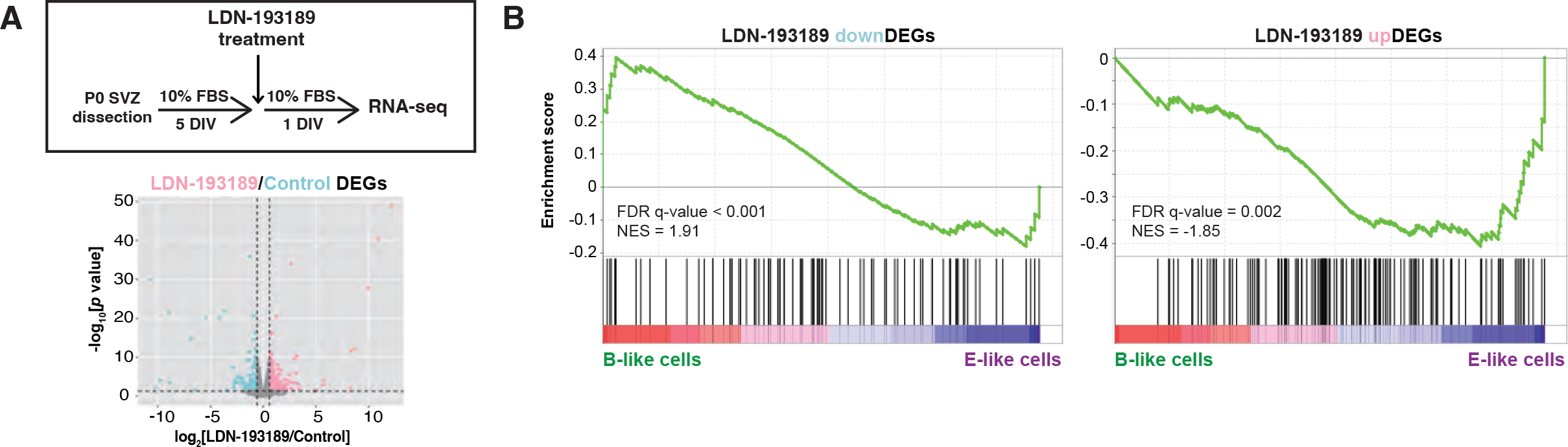
A BMP receptor inhibitor promotes aquisition of an E-like cell transcriptional profile in vitro (A) P0 NPC cultures maintained for 5 DIV with 10% FBS and then for 1 day with 10% FBS in the absence or presence of LDN-193189 (0.1 μM) were subjected to RNA-seq analysis (top). The Volcano plot shows the changes in gene expression induced by LDN-193189 treatment (bottom). DEGs upregulated by LDN-193189 treatment (edgeR: *p* < 0.05, log2[fold change] > 0.585) are shown in pink, whereas those downregulated by LDN-193189 (upregulated in the control) (edgeR: *p* < 0.05, log2[fold change] < –0.585) are shown in light blue. (B) GSEA for E16.5 B-like versus E-like cells and DEGs found to be downregulated or upregulated by LDN-193189 in (A). LDN-193189–downregulated DEGs are enriched in B-like cells, whereas LDN-193189–upregulated DEGs are enriched in E-like cells.

**Figure S5.**
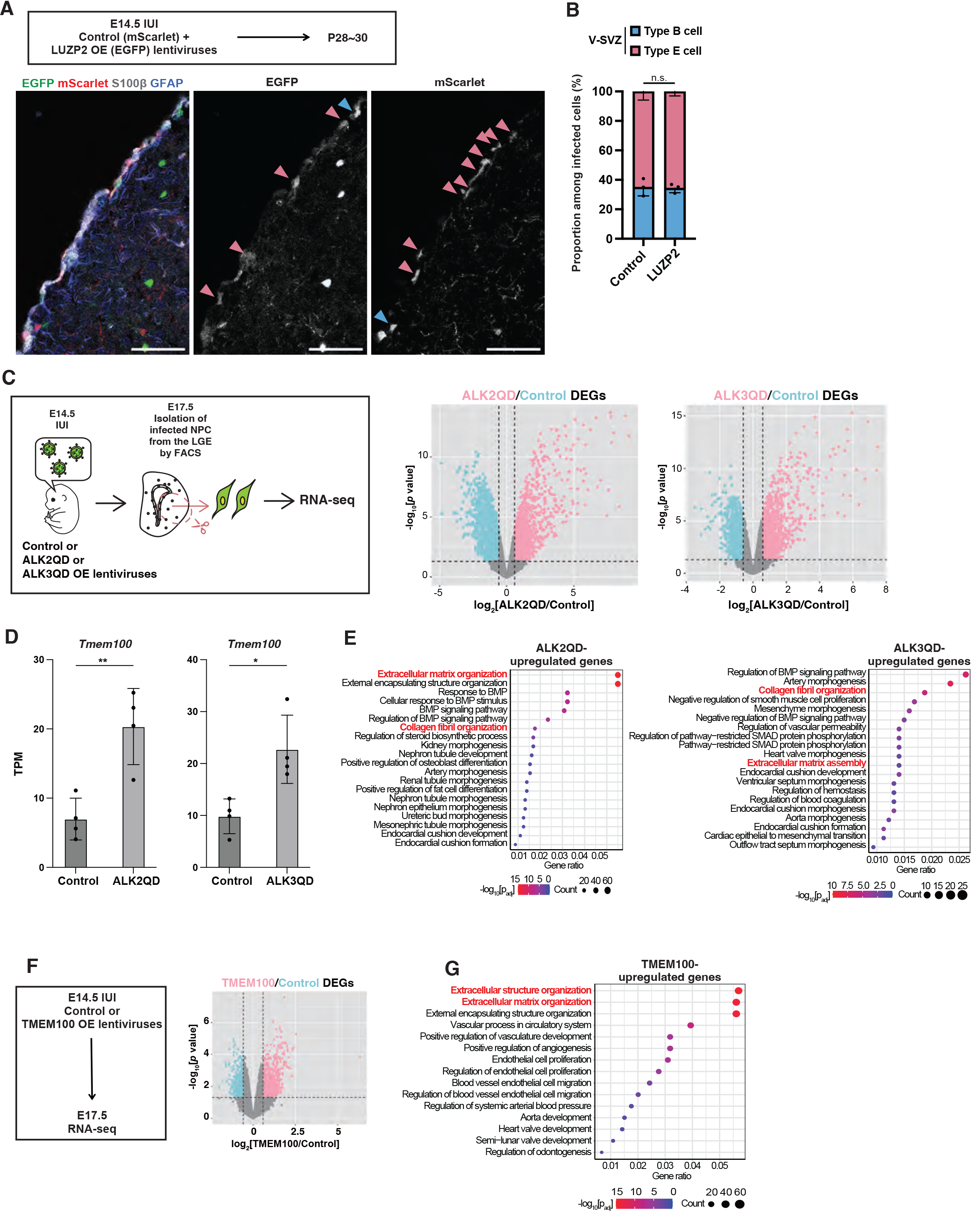
Analysis of the effects of B-like cell signature gene overexpression in vivo (A) Lentiviruses encoding mScarlet (FU-T2A-mScarlet-W) or both LUZP2 and EGFP (FU-LUZP2-T2A-EGFP-W) were injected into the embryonic lateral ventricle at E14.5, and brain sections of the resulting mice were subjected to immunohistofluorescence analysis of EGFP, mScarlet, GFAP, and S100β at P28–30. Light blue and pink arrowheads indicate young adult type B cells (defined as GFAP^+^ cells with apical attachment in the V-SVZ of the lateral wall) and type E cells (defined as S100β^+^ cells), respectively. Scale bars, 100 μm. (B) Proportions of young adult type B cells and type E cells among infected cells (control or LUZP2 overexpressing) in the V-SVZ determined from images as in (A). Cells positive for both mScarlet and EGFP were excluded from the analysis. Data are means ± SD (*n* = 3 mice). n.s., not significant (two-tailed Student’s *t* test). (C) RNA-seq analysis of EGFP^+^CD133^+^CD24^−^ NPCs isolated by FACS from the LGE of embryos at E17.5 that had been subjected to in utero infection at E14.5 with lentiviruses encoding EGFP alone (FU-T2A-EGFP-W) or both EGFP and either ALK2QD (FU-ALK2QD-T2A-EGFP-W) or ALK3QD (FU-ALK3QD-T2A-EGFP-W). The volcano plots show changes in gene expression induced by ALK2QD or ALK3QD expression. DEGs upregulated by ALK2QD or ALK3QD (edgeR: *p* < 0.05, log2[fold change] > 0.585) are shown in pink, whereas those downregulated by ALK2QD or ALK3QD (upregulated in the control) (edgeR: *p* < 0.05, log2[fold change] < –0.585) are shown in light blue. (D) Transcripts per million (TPM) values for *Tmem100* expression determined as in (C). Data are means ± S (*n* = 4 independent experiments). \**p* < 0.05, \*\**p* < 0.01 (two-tailed Student’s *t* test). (E) GO enrichment analysis for genes upregulated by ALK2QD or ALK3QD overexpression in (C). Extracellular matrix–related GO terms are highlighted in red. (F) RNA-seq analysis of EGFP^+^CD133^+^CD24^−^ NPCs isolated by FACS from the LGE of embryos at E17.5 that had been subjected to in utero infection at E14.5 with lentiviruses encoding EGFP alone (FU-T2A-EGFP-W) or both EGFP and TMEM100 (FU-TMEM100-T2A-EGFP-W). The volcano plot shows changes in gene expression induced by TMEM100 overexpression. DEGs upregulated by TMEM100 (edgeR: *p* < 0.05, log2[fold change] > 0.585) are shown in pink, whereas those downregulated by TMEM100 (upregulated in the control) (edgeR: *p* < 0.05, log2[fold change] < –0.585) are shown in light blue. (G) GO enrichment analysis for genes upregulated by TMEM100 overexpression in (F). Extracellular matrix–related GO terms are highlighted in red.

**Figure S6.**
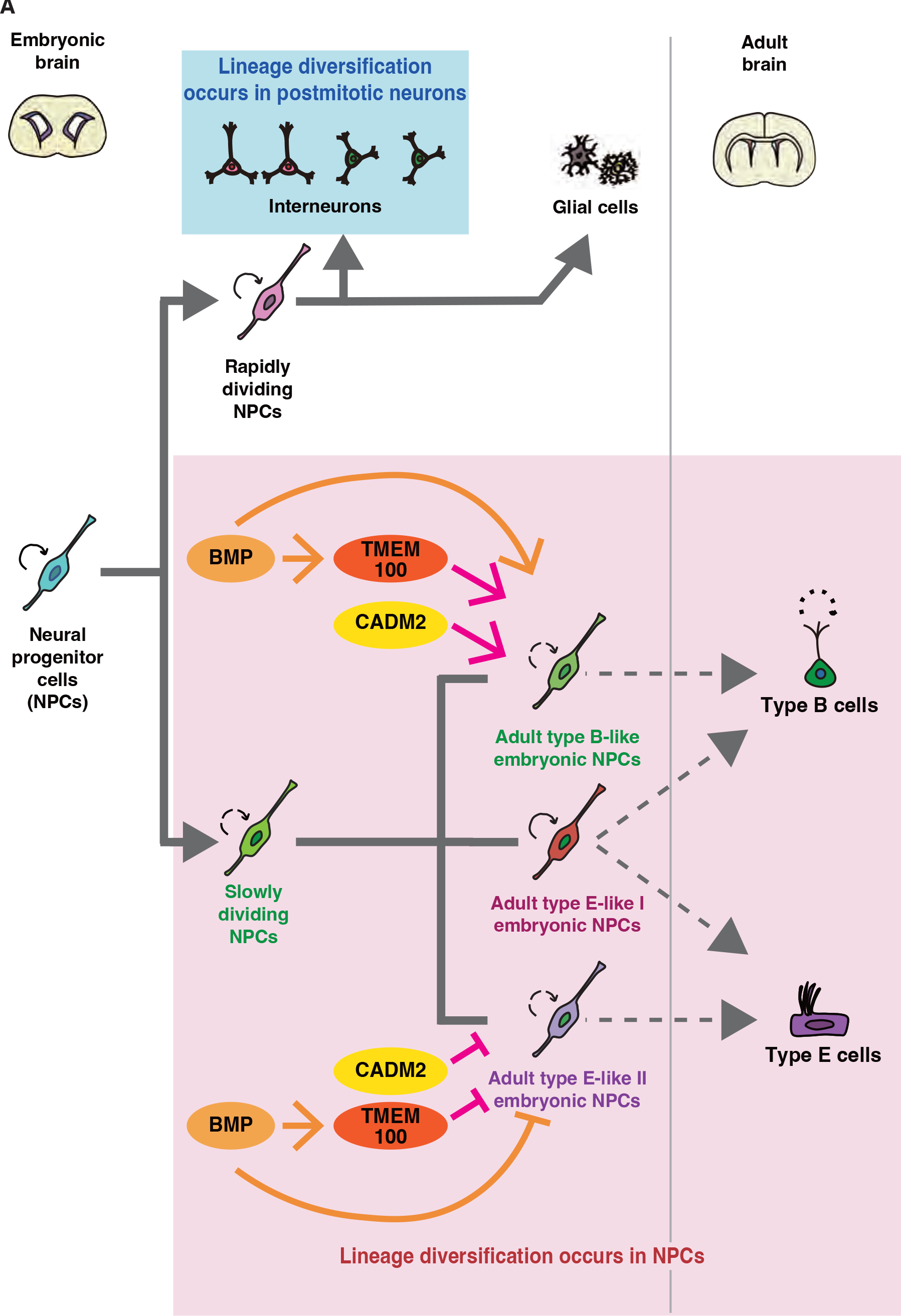
Model for the mechanisms underlying lineage diversification from embryonic NPCs to adult type B versus type E cells We previously showed that NPCs in the GE can be classified into at least two groups: rapidly dividing and slowly dividing cells ^9^. Rapidly dividing NPCs give rise to interneurons and glial cells, which then contribute to brain development. Lineage diversification among interneurons is thought to occur mostly among postmitotic cells (Bundler et al., 2022). In contrast, our present study suggests that lineage diversification between adult type B and type E cells occurs in NPCs. Slowly dividing NPCs thus comprise at least three distinct subsets: B-like cells, E-like I cells, and E-like II cells. B- like cells and E-like cells are predicted to give rise to adult type B cells and type E cells, respectively. Of note, the finding that E-like I cells possess characteristics of both adult type B cells and ependymal cells is consistent with the previous detection of common progenitors for these two adult cell types ^16,17^. With regard to the molecular mechanisms underlying lineage diversification between adult type B and type E cells, BMP signaling, CADM2, and TMEM100 contribute to the preferential production of adult type B cells rather than type E cells.

